# Proteostasis and Unfolded Protein Response Dynamics in Human Neuron and Mouse Glia Co-culture Reveal Cell-Specific Aging Responses

**DOI:** 10.1101/2025.08.11.669714

**Authors:** Lea A. Barny, Sarah K. Garcia, Aiden J. Houcek, Burak Uzay, Minsoo Kim, Ege T. Kavalali, Lars Plate

## Abstract

Proteostasis, or protein homeostasis, is a tightly regulated network of cellular pathways essential for maintaining proper protein folding, trafficking, and degradation. Neurons are particularly vulnerable to proteostasis collapse due to their post-mitotic and long-lived nature and thus represent a unique cell type to understand the dynamics of proteostasis throughout development, maturation, and aging. Here, we utilized a dual-species co-culture model of human excitatory neurons and mouse glia to investigate cell type– specific, age-related changes in the proteostasis network using data-independent acquisition (DIA) LC-MS/MS proteomics. We quantified branch-specific unfolded protein response (UPR) activation by monitoring curated effector proteins downstream of the ATF6, IRE1/XBP1s, and PERK pathways, enabling a comprehensive, unbiased evaluation of UPR dynamics during neuronal aging. Species-specific analysis revealed that aging neurons largely preserved proteostasis, although they showed some signs of collapse, primarily in ER-to-Golgi transport mechanisms. However, these changes were accompanied by upregulation of proteostasis-related machinery and activation of the ATF6 branch, as well as maintenance of the XBP1s and PERK branches of the UPR with age. In contrast, glia exhibited broad downregulation of proteostasis factors and UPR components, independent of neuronal presence. Furthermore, we quantified stimulus-specific modulation of select UPR branches in aged neurons exposed to pharmacologic ER stressors. These findings highlight distinct, cell-type-specific stress adaptations during aging and provide a valuable proteomic resource for dissecting proteostasis and UPR regulation in the aging brain.

**Significance:** Understanding how the unfolded protein response (UPR) and proteostasis network change with age is often studied in model organisms, where pathways are assessed across mixed cell types. Such systems can obscure cell-type-specific regulation. Here, we evaluate age-associated remodeling of the UPR and proteostasis network in a dual-species co-culture of human neurons and mouse glia using DIA proteomics. This approach enables species-specific proteomic profiling without physical separation, supported by a customizable data analysis pipeline. We show that neurons and glia exhibit divergent age-related responses, with neurons maintaining adaptive proteostasis and glia showing broader declines. The analytical framework presented here supports future studies to uncover additional cell-type-specific aging phenotypes or to probe the effects of pharmacologic or physical manipulation of biological systems.

## Introduction

Proteostasis, or protein homeostasis, is vital for maintaining neuronal health throughout aging^1^. Unlike many other cell types, neurons are post-mitotic and long-lived, making them particularly susceptible to the accumulation of misfolded or damaged proteins over time^2^. The proteostasis network contains molecular chaperones, the ubiquitin-proteasome system, autophagy, and stress signaling pathways such as the unfolded protein response (UPR) for regulation. A robust proteostasis network is essential for proper protein folding, trafficking, and degradation^3^. However, studies in *C. elegans*^4–7^, rodents^7–10^, and humans^11–13^ have shown that these systems decline with age, leading to protein aggregation and cellular stress that compromise synaptic function and neuronal viability^14,15^. Moreover, disrupted proteostasis is a hallmark of several age-related neurodegenerative diseases, including Alzheimer’s and Parkinson’s, underscoring its central role in preserving brain function^11^. Thus, elucidating the complex regulation of proteostasis networks and the UPR during human neuronal maturation and aging is critical for uncovering the mechanisms behind cognitive decline and identifying new therapeutic targets.

Previous work using excitatory human neurons revealed that endoplasmic reticulum (ER) stress induces UPR activation as a consequence of calcium dysregulation^16^. These findings were further supported by immunoblotting and fluorescence microscopy analyses of select chaperones, transcription factors, and the activation of the transmembrane receptors IRE1 and PERK, two conserved signaling branches of the UPR. Additionally, UPR activation was linked to impaired action potential-evoked synchronous release and a downregulation of key presynaptic proteins^16^. However, a comprehensive and unbiased characterization of this model is required to identify additional age-related disruptions in cellular pathways, particularly those involving the proteostasis network and broader remodeling of the UPR.

Thus, in the present study, we systematically profiled the contributions of each branch of the UPR during neuronal aging using data independent acquisition (DIA) liquid chromatography-mass spectrometry (LC-MS/MS) proteomics. To assess branch-specific UPR activation, we quantified the abundance of proteins previously defined by our group as selective effector targets of UPR transcription factors for each arm, then performed aggregate analyses to evaluate overall pathway engagement^17^. This method offers a high-throughput, quantitative, and unbiased approach to assess the state of the UPR across a broad range of samples. By enabling detection of multiple downstream targets, this method circumvents the need to directly identify transmembrane receptors or transcription factors, making it broadly applicable to global datasets during post-analysis without the need for specialized acquisition parameters. Additionally, data-independent acquisition (DIA) has been shown to capture a greater number of UPR-related proteomic targets than data-dependent acquisition (DDA)^17^.

The *in vitro* model of human neurons used here and numerous previous studies is comprised of neurons derived from H1 human embryonic stem cells (hESCs) and glial cells procured from mice^16,18–22^. This dual-species co-culture model enabled us to evaluate age-associated changes in proteostasis and other dysregulated biological processes in both neurons and glia and assess neuron-glia crosstalk using corresponding glial monocultures as a reference. Typically the combined cell types must be distinguished using techniques such as fluorescence-labeling (with proteins or dyes) followed by fluorescence-assisted cell sorting (FACS)^23^ or stable isotope labeling by amino acids in cell culture (SILAC)^24,25^. However, in dual-species models analyzed by LC-MS/MS, peptides can be confidently assigned to their species of origin based on sequence identification. This enables the detection of species-specific proteomic changes without the need for physical manipulation of the culture, an important advantage when working with sensitive systems like neurons, where such interventions could introduce experimental artifacts^26,27^. Similar co-culture systems have been previously utilized to study microbial interactions^28,29^, cancer^23,25,30^, and neuroscience^31^. DIA is ideal for analyzing complex mixtures commonly encountered in co-culture experiments because it offers enhanced and robust detection of low-abundance peptides across replicates despite differences in species-specific protein abundance^32^.

We found that neurons and glia exhibited distinct age-related proteomic responses. Proteostasis was largely preserved in aging neurons, with the exception of disruptions in ER-to-Golgi trafficking. In contrast, aging glia showed a diminished response across multiple cellular stress pathways and a marked reduction in core components of the proteostasis network, regardless of neuronal presence, indicating a broader age-associated decline in their homeostatic capacity. In terms of specific UPR regulation, aging neurons displayed significant upregulation of ATF6-associated markers, while XBP1s and PERK targets remained relatively stable in aged neurons compared to mature ones. Conversely, glial cells exhibited a broad downregulation of UPR associated proteins with age. Overall, at the evaluated timepoints, neurons exhibit a more adaptive proteostasis network during aging compared to glial cells. Our findings highlight a cell-type specific divergence in the proteostasis dynamics and UPR regulation and provide a valuable resource for understanding these dynamics during human neuronal aging.

## Results

### Development of a Species-Specific Peptide Detection Approach

We first established a pipeline to differentiate the proteome changes in aging human neurons from the glial feeder layer in the co-culture system. Human embryonic stem cells (hESCs) were transduced with the transcription factor neurogenin-2 (Ngn2) to induce synapse-forming cortical excitatory neurons (iN cells)^19^. These neurons were cultured on a glial feeder layer isolated from mouse hippocampal and cortical tissue, creating a dual-species co-culture model. iN cells were considered mature at 30 days and subsequently aged to 60 days^16^. Co-cultures and glia monocultures were collected at 30, 40, 50, and 60 days for LC-MS/MS analysis (**Fig. 1A**). To analyze the dual-species co-culture samples, an *in silico* tryptic digestion of the human and mouse proteomes, containing 20,434 and 17,214 entries, respectively, was performed in R using the *cleaver* package (**SI, Fig. S1A**). The resultant peptides were classified into six categories: human-only (proteotypic or razor), mouse-only (proteotypic or razor), and shared (proteotypic or razor). Approximately one-third of the theoretical peptides were determined to be either proteotypic human, proteotypic mouse, or shared (**SI, Fig. S1B**). LC-MS/MS data were analyzed using DIA-NN (v1.8.1), and the detected peptides were matched against a theoretical peptide database to determine species assignments, yielding three distinct data frames (**Fig. 1B**). This approach prevents species assignment of a given peptide based on the degree of protein level evidence (parsimony principle), which becomes increasingly complex when two species databases are being searched simultaneously^26,27^.

**Figure 1.**
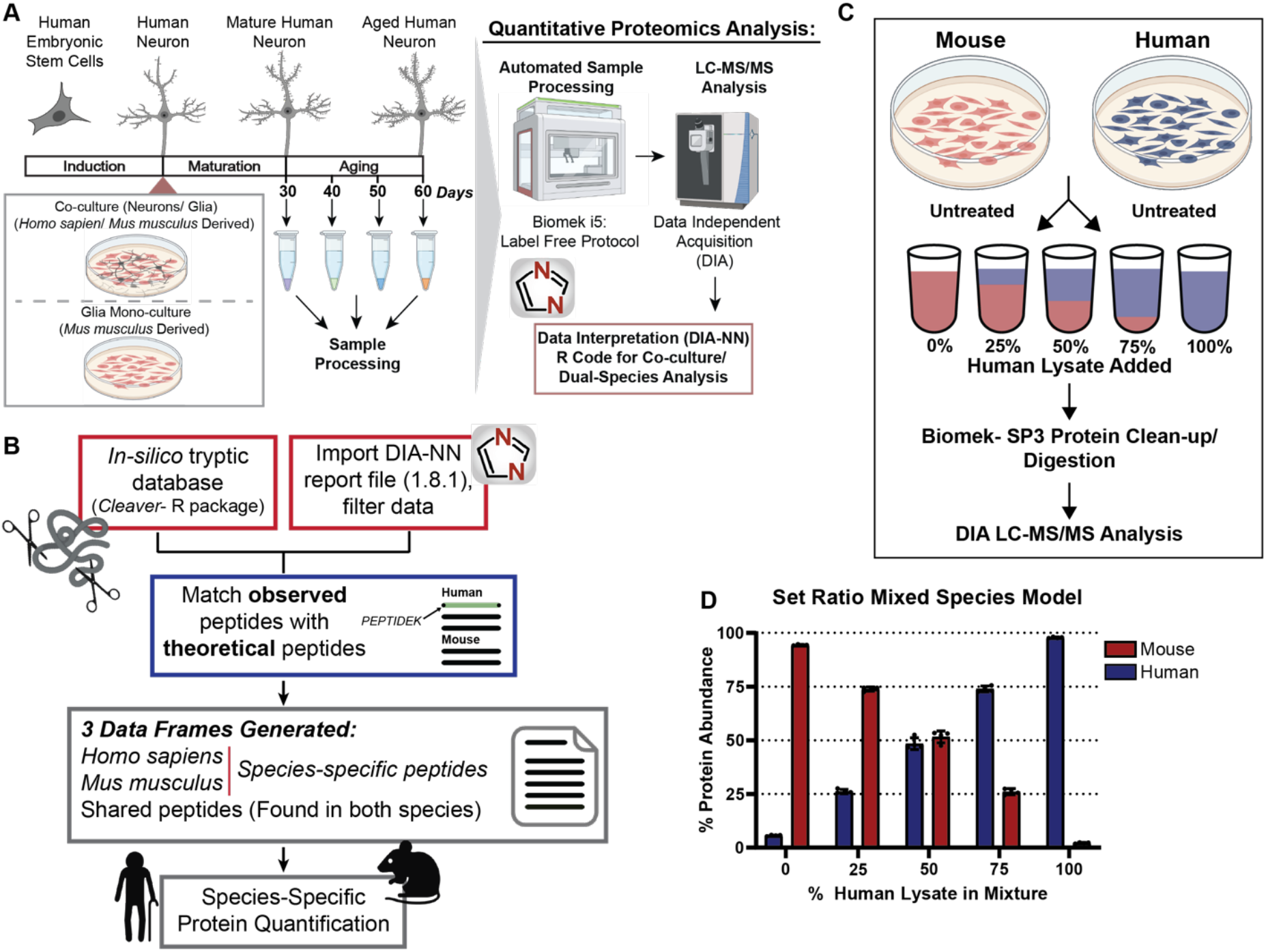
Development and validation of a pipeline to analyze dual-species co-culture samples via DIA LC-MS/MS. **(A)** Workflow for the collection and sample preparation of aged co-cultures (containing neurons (human derived) and glia (mouse derived)) and glia (mouse derived) monocultures for MS analysis via DIA (n = 10). **(B)** Overview of the data processing pipeline to achieve species-specific protein quantification from a DIA-NN (1.8.1) report file and our custom R code. **(C)** Experimental design of the mixed-species set ratio model system used to evaluate the performance of our computational pipeline. **(D)** Percent protein abundance of human and mouse proteins in the mixed species set ratio model (n = 4). Bars correspond to mean and error bars represent ± standard deviation (SD).

To validate our established computational pipeline, lysate of *M. musculus* N2a and *H. sapiens* SH-SY5Y cells, which are both of neuronal lineage, were mixed in set ratios (0%, 25%, 50% 75%, 100%) (**Fig. 1C**). Mixtures with 25-75% SH-SY5Y lysate were determined to have a similar number of human-unique protein identifications despite the decrease in amount of human material on-column, indicating peptide detection remains stable over a range of concentrations using our DIA method (**SI, Fig. S1C**). Additionally, using our mixed-species set ratio samples, we quantified the false positive rate of peptides of the opposite species after applying different filters to the DIA-NN output file (**Dataset S1**). Given that human peptides and proteins should be absent in the 100% *M. musculus* lysate sample, and vice versa, this method enabled us to assess the accuracy of peptide species-assignment and the impact of filtering strategies at the peptide and protein level. The addition of a precursor posterior error probability (PEP) score cutoff of 0.01 resulted in a significant decrease in the false positive rate of human protein identifications in the 100% *M. musculus* lysate by approximately 4% (**SI, Fig. S2A**). Similarly, the mouse protein identifications in the 100% *H. sapiens* lysate samples decreased by 4% using the PEP score to increase filtering stringency (**SI. Fig. S2B**). However, the percentage of human false positive proteins was greater than mouse false positive proteins in their respective single species samples with the PEP filter, 6.26 ± 0.22% and 2.99 ± 0.15%, respectively. To avoid misassignment of shared peptide quantifications, peptides with ambiguous species origin were excluded from the analysis. Overall, the applied peptide filtering criteria produced human and mouse protein abundance profiles that closely reflected their respective concentrations, demonstrating that our data analysis pipeline can accurately distinguish peptides from two species (**Fig. 1D**).

To assess whether the database size in DIA-NN affects the number of false positive identifications, we searched 100% human and mouse samples using three different approaches: (1) their respective species-specific FASTA, (2) the FASTA of the opposite species, and (3) a combined human and mouse FASTA. DIA-NN searches of the 100% mouse lysate using a human FASTA database, and vice versa, yielded a higher number of false positives compared to searches using a combined mouse-human FASTA (**SI, Fig. S3**). Thus, to prevent inflation of false positive identifications, we used a combined mouse-human FASTA to analyze all dual species samples in DIA-NN. Lastly, to determine if peptide and protein identifications could be improved for dual species samples in DIA-NN with the use of a sample-specific spectral library, mixed species set ratio samples were fractionated with gas phase fractionation (GPF)-DIA to make a spectral library in DIA-NN. Briefly, GPF enables the use of smaller isolation windows (i.e. 2-4 *m/z*) before peptide fragmentation by injecting the sample multiple times across a defined mass range (i.e. 400-1,000 *m/z*). This narrower isolation window improves precursor selection specificity, reducing peptide co-fragmentation and minimizing interference from overlapping precursors and fragments, ultimately enhancing data interpretation^36,42^. However, the use of GPF-DIA to build a sample specific library did not result in a greater number of peptide or protein identifications in DIA-NN when searching the mixed species set ratio dataset (**SI, Fig. S4; Dataset S2**). Thus, all samples throughout the study were analyzed with library-free searching in DIA-NN.

### Distinct Aging Trends Between Neurons and Glia

To achieve species-specific protein quantification, neuronal and glial peptide intensities from the day 30 to day 60 aging experiment (**Fig. 1A**) were median-normalized, and protein-level quantification was performed using the MaxLFQ^38^ function in R (**SI Fig. S5 & S6; Dataset S3**). Glial proteins were identified at approximately twice the frequency of neuronal proteins in the co-culture samples, consistent with the higher initial glial seeding density compared to neurons. Principal component analysis (PCA) of aged neurons and glia in co-culture samples revealed age-dependent clustering, indicating a robust impact of aging on proteomic profiles (**Fig. 2A**). Overall, neurons showed a greater number of significantly upregulated proteins with age, whereas glia predominantly exhibited downregulated differentially expressed proteins (**Fig. 2B**). Both neurons and glia displayed the highest number of differentially expressed proteins at the day 60 timepoint. Minimal overlap was observed in significantly upregulated and downregulated proteins across all three timepoints in neurons. However, both neurons and glia at days 50 and 60 showed a greater number of shared proteins, with day 60 exhibiting the highest number of uniquely downregulated proteins in both cell types (**Fig. 2C; SI, Fig. S7A-B**). To investigate the biological processes associated with the upregulated and downregulated proteins at day 60 in neurons, gene ontology analysis was carried out using Enrichr^39–41^. Oxidative phosphorylation and processes related to mitochondrial function such as translation, membrane organization, and cristae formation were significantly upregulated in neurons with age (**Fig. 2D**). Numerous other cellular processes were also upregulated with age in neurons, including response to unfolded protein and ER stress, synapse organization, and cellular response to oxidative stress and superoxide. Downregulated cellular processes included mRNA splicing and processing, telomere maintenance and organization, and DNA damage repair (**Fig. 2E**). Several processes related to proteostasis were also downregulated with age, including those involved in protein trafficking from the ER to the Golgi apparatus, such as machinery involved in COPII coated vesicle budding and cargo loading, as well as SRP dependent co-translational targeting of proteins to the ER membrane. Notably, all four isoforms of SEC24 (A, B, C, and D), key components of the COPII complex required for the export of proteins from the ER to the Golgi, were significantly downregulated with age in human neurons (**SI, Fig. S8**).

**Figure 2.**
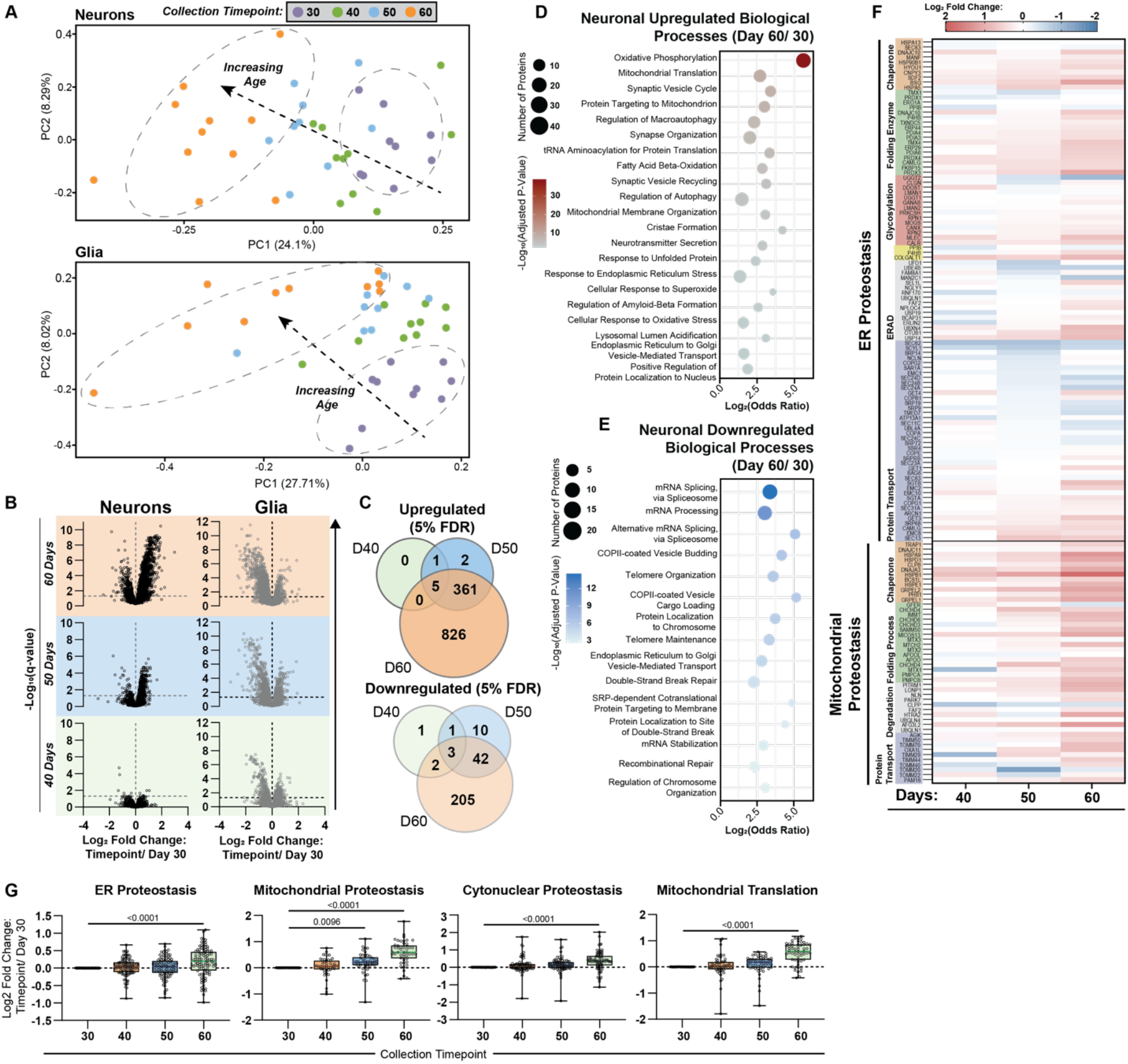
Global proteome changes in co-culture neurons and glia with age. **(A)** Principal component analysis (PCA) plots of co-culture neurons and glia at days 30 (purple), 40 (orange), 50 (blue), and 60 (green). **(B)** Volcano plots depicting aged cocultures, categorized by neurons and glia from human and mouse species, respectively. Aged cocultures at 40, 50, and 60 days were compared to the control condition at Day 30. Proteins with a −log_10_(q value) ≥ 1.3 (5% FDR) were considered significant, indicated by a horizontal dashed line. **(C)** Significantly upregulated and downregulated proteins (5% FDR) shared and unique to the 40 (orange), 50 (blue), and 60 (green) day collection timepoints. Gene ontology of the significantly upregulated **(D)** and downregulated (**E**) proteins at 60 days using Enrichr^1^. Adjusted p-value was computed using the Benjamini-Hochberg method to correct multiple hypothesis testing. **(F)** Heatmap of log_2_ fold change of neuronal proteins involved in ER and mitochondrial proteostasis, identified at days 40, 50, and 60 relative to the day 30 control. **(G)** Boxplots showing the log_2_ fold change (collection timepoint/day 30) of neuronal proteins involved in ER function, mitochondrial activity, cytonuclear proteostasis, and mitochondrial translation, plotted collectively. n = 10; data are presented as mean ± SEM. Statistical analysis was performed using one-way ANOVA, followed by Dunnett’s post hoc test for multiple comparisons against the day 30 control. *p* < 0.05 was considered statistically significant.

Given the striking changes to proteostasis terms identified in this unbiased pathway analysis, we decided to further investigate how the proteostasis network is altered in human neurons. We created a heatmap of key ER and mitochondrial proteostasis markers, established by the Proteostasis Consortium^43,44^ (**Fig. 2F**). ER-resident folding enzymes and chaperones were largely upregulated with age, reaching peak abundance at day 60. In contrast, many components of the ER protein transport machinery were progressively downregulated over time. Several aspects of mitochondrial proteostasis, including chaperones, protein folding and degradation systems, as well as protein import machinery, were also consistently upregulated with age. Statistical analysis of the log_2_ fold change showed that terms related to different aspects of proteostasis, including ER, mitochondrial and cytonuclear proteostasis, were significantly upregulated at 60 days compared to 30 days (**Fig. 2G**). Cytosolic and mitochondrial translation were significantly upregulated with age by day 60 (**Fig. 2G, SI, Fig. S9A-B**). In addition, degradation pathways such as the ubiquitin proteasome system and the autophagy lysosome pathway showed increased activity as a function of age, with the autophagy lysosome pathway becoming significantly upregulated starting at day 50. Thus, the components of the proteostasis network in neurons were largely upregulated with age, pointing to neurons retaining the ability to induce these components in response to aging stimuli.

In contrast, glial cells displayed a markedly distinct proteomic profile compared to neurons. Many biological processes that were upregulated in neurons were downregulated in glia, including pathways associated with the ER stress response, cellular response to oxidative stress, mitochondrial translation and organization, and tRNA aminoacylation for protein synthesis at 60 days compared to 30 days. ER-to-Golgi vesicle-mediated transport was downregulated in both glia and neurons, suggesting a shared suppression of protein trafficking. Notably, glial cells also showed downregulation of pathways involved in sterol and UDP-N-acetylglucosamine biosynthesis, sphingolipid catabolism, and protein degradation (i.e., ER-associated degradation, ubiquitin-dependent protein catabolism, regulation of autophagy). Additionally, immune-related processes were diminished in glia (i.e., Toll-like receptor signaling, regulation of type I interferon production, regulation of interferon-beta production, and cellular response to type II interferon) (**SI. Fig. S10A**). Conversely, cell respiration, specifically oxidative phosphorylation, was upregulated in aging glia at day 50 compared to day 30, although the increase was not as pronounced as in neurons. Fatty acid oxidation, glutathione, cholesterol, and serine metabolic processes were also upregulated with age (**SI. Fig. S10B**). In summary, neurons exhibited relatively mild signs of proteostasis collapse compared to glia, particularly associated with the downregulation of ER trafficking-related components. Neurons appear to retain the capacity to activate compensatory mechanisms to restore proteostasis. In contrast, glial cells display widespread downregulation of critical components of the proteostasis network.

### UPR Activation by LC-MS/MS in a Highly Homologous Dual-Species Model

The UPR consists of three branches, named after their respective sensor proteins: ATF6 (Activating Transcription Factor 6), IRE1 (Inositol Requiring Enzyme 1), and PERK (Protein Kinase RNA-like ER Kinase). Activation of each arm results in the production of distinct transcription factors that transcriptionally and translationally remodel ER proteostasis pathways to combat ER stress (**Fig. 3A)**. We first validated that cell-type specific UPR activation can be effectively distinguished in the dual-species co-culture model. To do this, human SH-SY5Y cells were treated with ER stressors (tunicamycin Tm^45^; or thapsigargin Tg^46^), while mouse N2a cells were left untreated. SH-SY5Y lysate activated with either vehicle or global UPR activators (Tm or Tg) were mixed in a 1:4 ratio with N2a untreated lysate (**Fig. 3B**). Detection of UPR activation was determined in our LC-MS/MS datasets using branch-specific UPR proteomics targets previously elucidated by our group^17^. Activation of select UPR markers in vehicle, Tm, and Tg-treated SH-SY5Y cells was validated using quantitative western blot analysis (**SI, Fig. S11**).

**Figure 3.**
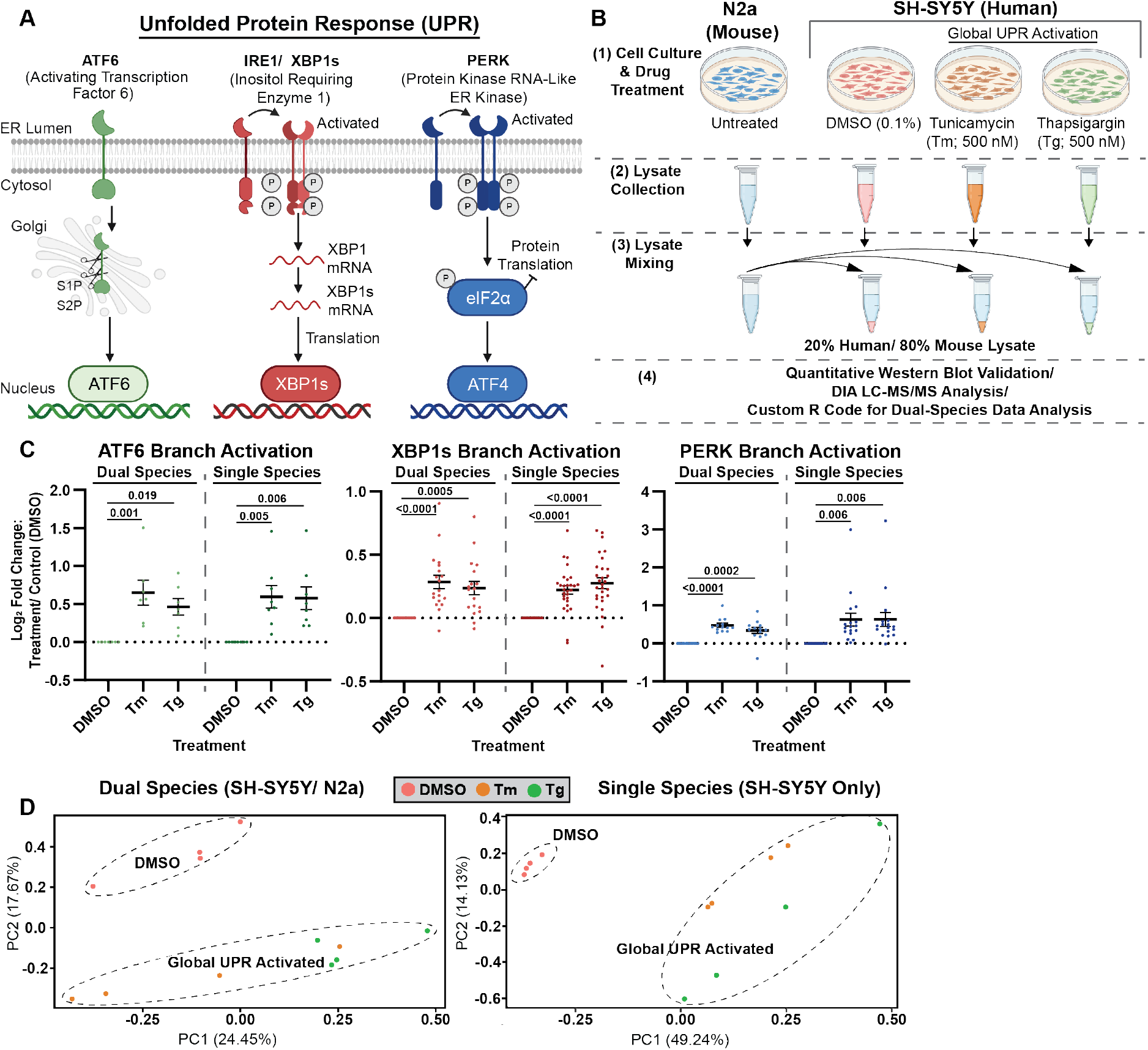
Detection of UPR activation in a mixed species set ratio SH-SY5Y/ N2a model. **(A)** Schematic showing the three major branches of the UPR (ATF6, XBP1s, and PERK) along with their respective transcription factors. **(B)** Samples generated to evaluate UPR activation in a mixed species versus single species model. SH-SY5Y cells were treated with DMSO (vehicle, control), Tm (500 nM), and Tg (500 nM) for 16 hours and lysate was collected. Additionally, untreated N2a cell lysate was collected. Human treated lysates (SH-SY5Y) were then mixed with mouse (N2a lysate) at a 1:4 ratio. SH-SY5Y single species UPR activated lysate was also analyzed as a control. **(C)** The log_2_ fold change of branch-specific UPR markers, previously elucidated by our group, at different timepoints in co-culture neurons and glia plotted collectively to determine branch-specific activation overtime. Dots represent individual proteins. Error bars represent the mean ± SEM. **(D)** PCA analysis to confirm separation of untreated (DMSO) and UPR activated (Tm/ Tg treated) in dual species (20% human and 80% mouse lysate) and single species (human) samples.

Of the 60 UPR protein activation markers, 59 were detected in single-species injections, while only 40 were identified in the dual-species (mixed) model (**Dataset S4)**. LC-MS/MS analysis of human SH-SY5Y cells in both single and mixed-species samples revealed significant activation of all three UPR branches following treatment with either Tm or Tg (**Fig. 3C**). PCA analysis further demonstrated a clear separation between DMSO-treated (control) and globally UPR-activated (Tm or Tg) human SH-SY5Y cells in both single- and dual-species samples (**Fig. 3D**). Together, these results validate our ability to detect UPR activation in a dual-species co-culture model despite the slightly lower identification of UPR markers.

### Age-Dependent Divergence in UPR Activation between human neurons and glia

Next, we evaluated the activation of the three UPR branches throughout the day 30 to day 60 aging time course in neurons and glia using the previously described branch-specific protein markers^17^. Neurons generally showed an increase in UPR activation with age (**Fig. 4A**), whereas glia exhibited an overall decrease in UPR activation with age (**Fig. 4B**). In neurons, ATF6 protein markers were significantly upregulated at day 60, while those of XBP1s and PERK showed a trending but non-significant increase over time. UPR activation was significantly downregulated at day 60 compared to day 30 across all three branches in glia. Additionally, XBP1s and PERK markers were significantly downregulated at day 50 compared to day 30.

**Figure 4.**
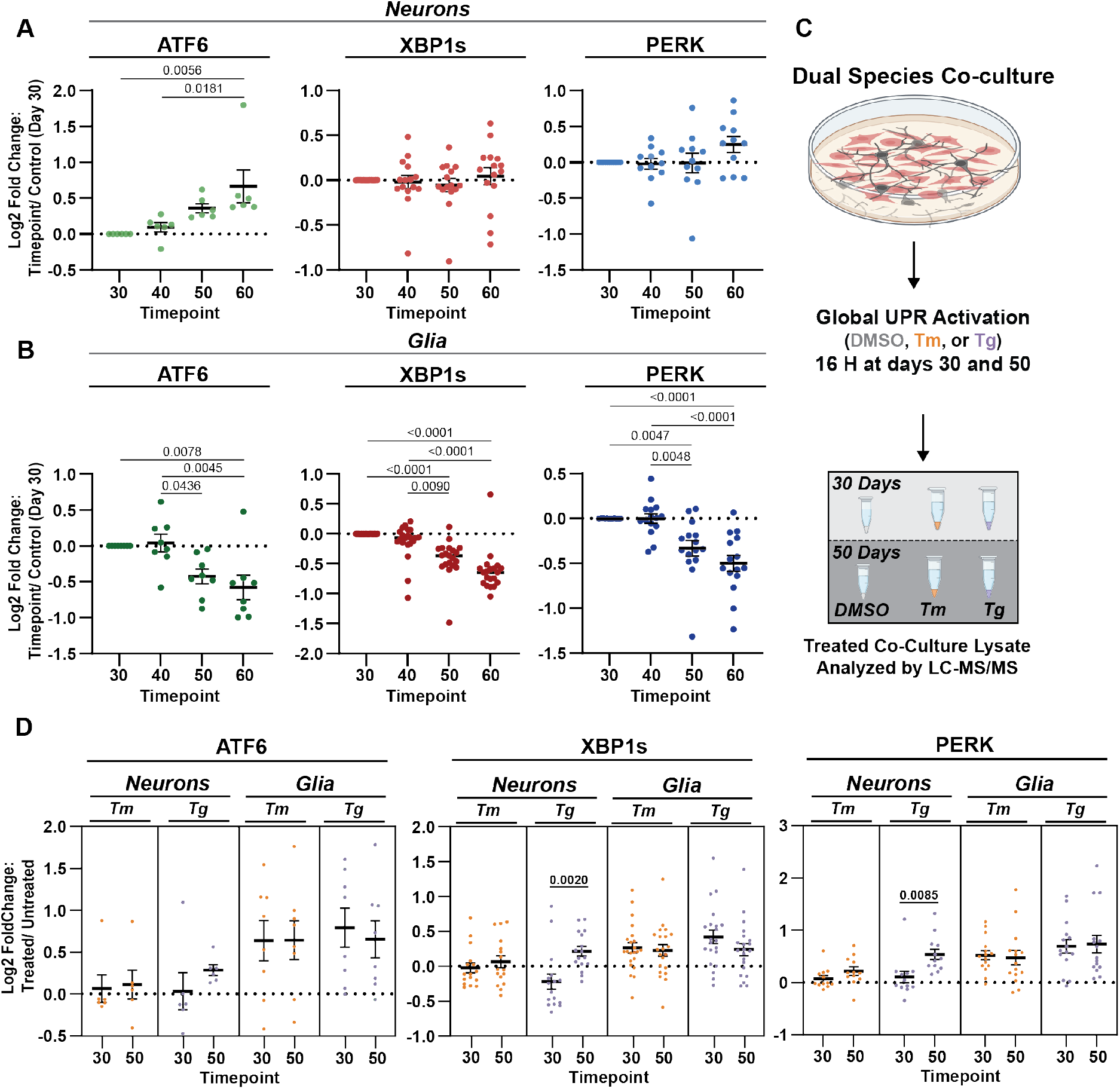
Quantification of branch-specific UPR markers in aging co-cultures. **(A)** The log_2_ fold change of UPR markers overtime in co-culture **(A)** neurons and **(B)** glia plotted collectively to determine branch-specific activation overtime. Branch specific UPR markers were previously elucidated^2^. Dots represent individual proteins. Error bars represent the mean ± SEM. **(C)** Experimental design, treatment of co-culture samples at days 30 and 50 with Tm (5 µM), Tg (300 nM), or DMSO for 16 hours. **(D)** log_2_ fold change of UPR markers in co-cultures treated with either Tm or Tg at days 30 or 50 compared to their untreated controls.

To assess whether age influences the capacity of neurons and glia to mount a UPR in response to ER stress, we acutely treated co-cultures with Tm (5 µM) or Tg (300 nM) at days 30 and 50 for 16 hours (**Fig. 4C; Dataset S5**). At day 30, neurons exhibited non-significant UPR activation following either treatment, while glia showed a robust UPR response to both Tg and Tm (**SI, Fig. S12A**). By day 50, neurons treated with Tg displayed significant activation of the XBP1s and PERK branches, with a small but non-significant increase in ATF6 activity (**Fig. 4D**). In contrast, Tm-treated neurons showed only marginal, non-significant increases across all three UPR branches at day 50 relative to day 30 (**SI, Fig. S12B**). Glia, however, maintained consistent UPR activation at both time points following acute treatment with either compound (**Fig. 4D**). In summary, glia exhibited distinct responses to UPR activators with age relative to neurons, which further show differential sensitivity to Tg versus Tm.

### Neuronal influence of glial cell proteomic profiles

To assess how neuronal presence alters glial behavior, we measured glia peptide intensities in monoculture and compared them to the neuronal co-cultured glia. Peptides were median-normalized, and protein quantification was performed using the MaxLFQ^38^ function in R (**SI, Fig. S13; Dataset S6**). Similar to co-culture glia, the more proteins in the monoculture glia were significantly downregulated than upregulated at later time points (days 50 and 60) compared to the 30-day control (**Fig. 5A**). All three branches of the UPR were also markedly suppressed in monoculture glia at days 50 and 60, mirroring the pattern observed in co-culture (**Fig. 5B**). When we compared the significantly downregulated proteins present in the co-culture and monoculture glia, we found 1,407 proteins in common, 383 and 1,540 proteins uniquely downregulated in the co-culture and monoculture, respectively (**Fig. 5C**). Moreover, the shared proteins exhibited a greater magnitude of downregulation in the co-culture condition than in monoculture (**Fig. 5D**). Overall, these data show that co-culture and monoculture glia are phenotypically similar with age; however, glia in co-culture exhibit greater downregulation of these proteins, which may reflect their supportive role in maintaining neuronal function.

**Figure 5.**
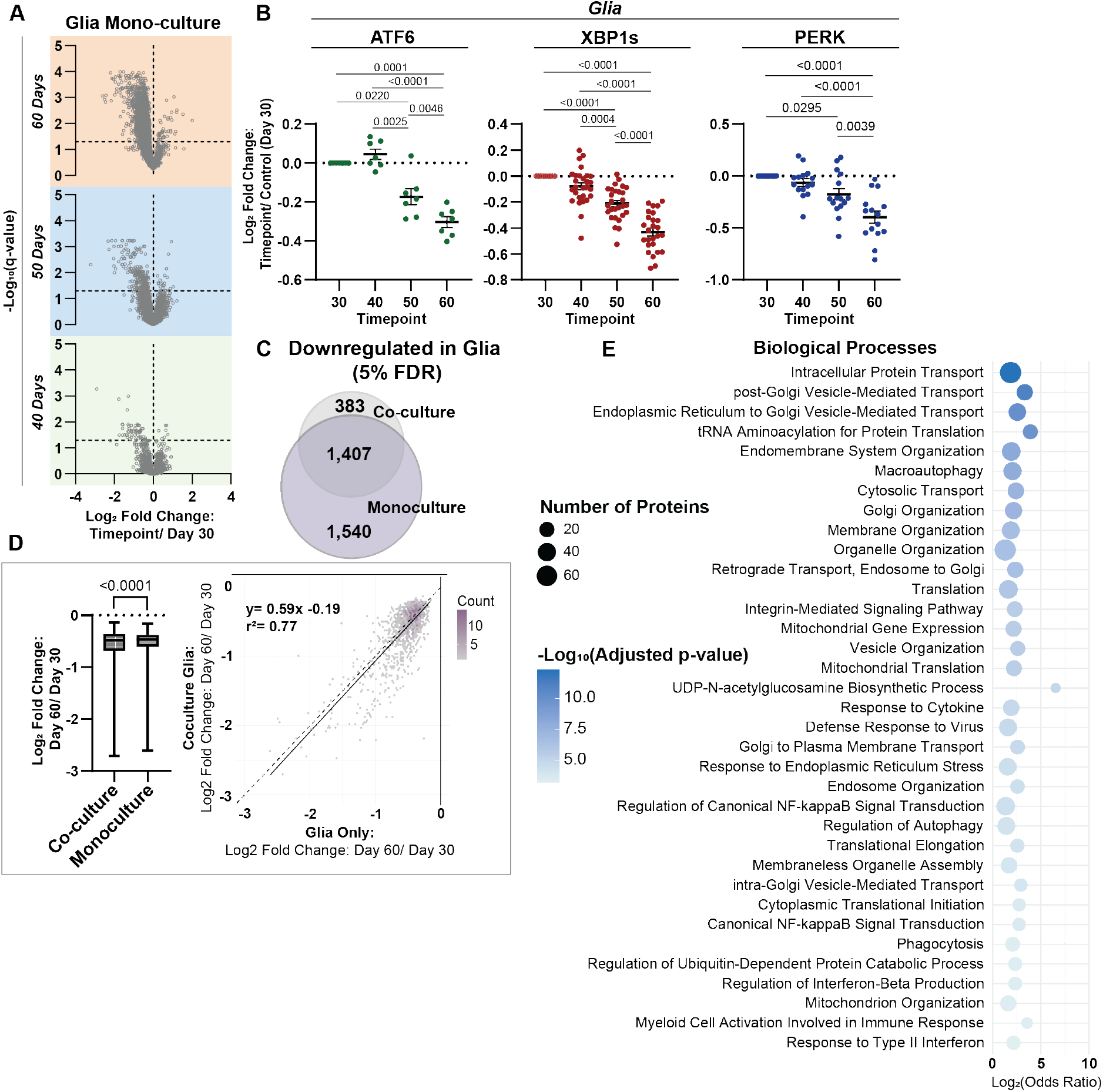
Comparing mono-culture and co-culture glia during aging. **(A)** Volcano plots of aged glia monocultures (40, 50 and 60 days) compared to the control (Day 30). Proteins with a −log_10_(q value) ≥ 1.3 (5% FDR) were considered significant. **(B)** The log_2_ fold change of UPR markers overtime in monoculture glia plotted collectively to determine branch-specific activation overtime. Error bars represent the mean ± SEM (n = 5). **(C)** Significantly downregulated proteins (5% FDR) shared and unique to the co-culture and monoculture glia at the 60 day timepoint. (**D**) Box and whisker plot to determine if there is a difference between the overall log fold changes (Day 60/ Day 30) of downregulated proteins shared between the co-culture and monoculture glia. Principal component analysis (PCA) plots of the log_2_ fold change of downregulated proteins (Day 60/ Day 30) in co-culture and monoculture glia depicted in the box and whisker plot. **(E)** Gene ontology of the significantly downregulated proteins shared between the co-culture and monoculture glia at day 60 using Enrichr^1^. Adjusted p-value was computed using the Benjamini-Hochberg method to correct multiple hypothesis testing.

To further explore the functional relevance of the shared downregulated proteins in monoculture and co-culture glia, we performed gene ontology (GO) analysis with Enrichr to identify associated biological processes (**Fig. 5E**). The most significantly downregulated processes included several pathways related to Golgi trafficking and organization such as post-Golgi vesicle-mediated transport, ER to Golgi vesicle-mediated transport, endosome to Golgi retrograde transport, and intra-Golgi vesicle-mediated transport. Additional downregulated processes involved mitochondrial translation and organization, tRNA aminoacylation for protein synthesis, response to ER stress, and the UDP-N-acetylglucosamine biosynthetic process. Immune-related processes were also affected, including responses to cytokines and viral infection, canonical NF-κB signaling, regulation of interferon-beta production, and response to type II interferon. Proteins exclusively downregulated in co-culture glia were implicated in biological processes such as astrocyte activation, astrocyte development, and mitochondrial translation (**SI, Fig. S14**). Together, these findings suggest that while aging independently suppresses a broad array of critical cellular functions in glia, the presence of neurons in co-culture further amplifies these changes in a subset of shared molecular pathways.

## Discussion

In this study, we developed a DIA-MS analysis pipeline to investigate dual-species co-cultures. We applied this workflow to characterize global proteomic changes in human neurons and mouse glia. We optimized statistical filtering parameters for peptides identified by DIA-NN using a dual-species set ratio model. This model confirmed that the measured protein abundances reliably reflected the true proportions of human and mouse proteins in mixed samples. Notably, human proteins remained consistently detectable even when their concentration was reduced to just 25% (125 ng of human peptides on-column), highlighting the high sensitivity of DIA-MS for this application. To address potential ion interference due to sample complexity and improve proteome coverage, we performed gas-phase fractionation (GPF) on the dual-species set ratio samples to generate a sample-specific spectral library^36^. Interestingly, this approach did not significantly increase the number of protein identifications. This likely reflects the effectiveness of DIA-NN’s deep learning-based spectral library prediction, which leverages machine learning to accurately model peptide fragmentation and retention times directly from the FASTA database. These *in-silico* predictions appear to capture a broad and representative peptide space, making empirical library generation unnecessary. Overall, this pipeline provides a flexible and scalable approach for analyzing complex co-culture systems and can be readily adapted to a wide range of experimental designs.

Ontology analysis revealed that human neurons exhibit an age-associated decline in telomere organization, maintenance, and DNA repair capacity. Interestingly, this decline was accompanied by an upregulation of oxidative phosphorylation and mitochondrial translation machinery, suggesting a compensatory increase in energy production pathways to support ongoing repair processes and stress responses, consistent with previous reports^47^. Alternatively, the observed increase in oxidative phosphorylation may result from elevated cytosolic calcium. Prior work by Uzay et al., using the same model system, demonstrated an age-related increase in N-methyl-D-aspartate receptor (NMDAR) expression, a key mediator of calcium influx in neurons^16^. This NMDAR overactivation was linked to an imbalance in excitatory synaptic transmission (E/I imbalance), a hallmark of aging and neurodegeneration^48,49^. Excessive NMDA receptor (NMDAR) activity can also disrupt calcium homeostasis, where excess calcium is redistributed to both the ER and mitochondria to buffer the overload^50,51^. Calcium influx into mitochondria is known to enhance oxidative phosphorylation and mitochondrial translation, but also contributes to increased oxidative stress^52,53^. Like neurons, glia maintained ATP production through active metabolic processes. However, ontology analysis revealed a more substantial decline in proteostasis in glia than neurons. Proteostasis impairments included ER-to-Golgi transport, ER-associated degradation (ERAD), autophagy, ER stress responses, and mechanisms that mitigate oxidative damage. In contrast, while glia retained machinery to remain metabolically active, they exhibited deficiencies in mounting adaptive or inflammatory responses. This suggests that proteostasis impairment in glia may precede or hinder immune activation, potentially contributing to a maladaptive cellular environment during aging. However, the lack of immune activation may be beneficial in preserving both glial and neuronal viability, as sustained neuroinflammation is known to contribute to cellular dysfunction and death^54^. Alternatively, the absence of astrocyte activation may suggest that these cells are not experiencing the anticipated level of stress associated with proteostasis decline, as proteostasis decline has been shown to result in glial activation^55,56^, potentially leading to a downregulation of UPR and other ER stress pathways.

In co-cultured excitatory neurons, we observed a significant age-dependent increase in ATF6-regulated protein activation as a function of age. In contrast, the XBP1s and PERK pathways showed non-significant increases in their associated translational markers at the same time point. While previous studies have linked aging to a decline in proteostasis, our findings suggest a more modest, potentially adaptive UPR response in this excitatory neuron model. As discussed above, excessive NMDAR activity can disrupt intracellular calcium homeostasis and compromise ER function, leading to the accumulation of misfolded and aggregated proteins. Consistent with this mechanism, previous studies have shown that treatment of cultured rat hippocampal neurons with NMDA to induce excitotoxicity activates the PERK arm of the UPR (with data available for this branch only), as demonstrated by western blot analysis^34^. In our co-culture system, fluorescence microscopy revealed increased expression of ER stress markers BiP and CHOP between days 30 and 50^16^, further supporting a connection between calcium dysregulation and ER stress in aging excitatory neurons.

Overall, while decreases in ER calcium levels have been extensively studied in neurons and other model organisms using the pharmacological inhibitor Tg^5,34,57^, the effects of ER calcium overload remain poorly understood. Interestingly, a study in TMCO1-deficient HeLa cells, where TMCO1 serves as a critical ER membrane calcium leak channel, demonstrated that ER calcium overload can activate the IRE1/XBP1s branch of the UPR, promoting cell survival^58^. Conversely, in *C. elegans*, inhibition of ATF6 has been shown to modulate calcium homeostasis and mitochondrial signaling, resulting in an increased lifespan, highlighting a complex relationship between calcium dynamics, stress signaling, and longevity^59^. Future studies will be needed to determine the functional relevance of ATF6 activation in this context as well as whether it contributes to neuron survival. It is also possible that ATF6 pathway activation in neurons may result from impaired COPII-mediated trafficking, which facilitates protein transport from the ER to the Golgi apparatus. In general, disruptions in ER-to-Golgi trafficking are known to trigger ER stress and activate the UPR in neurons ^60,61^. In human neurons, we observed a significant decrease in the expression of SEC24 isoforms over time, which are essential components of the inner coat protein complex II (COPII) machinery. These isoforms directly interact with cargo proteins or adaptors to mediate proper sorting into COPII vesicles^62,63^. Previous studies have demonstrated that postmitotic neurons are particularly sensitive to the loss of SEC24C, which leads to UPR activation and apoptosis—an effect not rescued by the overexpression of SEC24D^60^. Consistent with these findings, we observed reduced expression of both SEC24C and SEC24D in human neurons, suggesting that impaired trafficking may contribute to the selective activation of the ATF6 pathway and possibly impact neuronal viability during aging.

We observed a significant age-dependent decline in the expression of proteins regulated by all three branches of the UPR by 60 days in glia. Notably, a significant reduction was also evident at day 50 in the XBP1s and PERK arms of the UPR. These findings suggest that either the UPR receptors themselves (ATF6, XBP1s, and PERK) or downstream transcription factors may be downregulated or mislocalized^5,64^ with age, associated genes are improperly transcriptionally regulated^64^ or that epigenetic mechanisms^65,66^ may be contributing to this decline. In support of this, previous work has shown that the capacity to induce expression of key UPR genes, such as XBP1s, HSPA5, and CHOP, is significantly reduced in the hippocampus of middle-aged and aged mice compared to young animals following experimental induction of ER stress^67^. However, the cell-type specificity of this diminished response was not addressed in those studies. Our findings suggest that age-related repression of the UPR occurs predominantly in glia, whereas neuronal UPR signaling remains preserved, indicating neurons employ distinct mechanisms to retain UPR signaling with age.

To investigate cell-type-specific UPR responsiveness with age, we challenged co-cultures with global UPR activators, tunicamycin (Tm) and thapsigargin (Tg), at days 30 and 50. Glia exhibited a comparable UPR response at both time points. In contrast, neurons mounted a stronger response at day 50 than at day 30, specifically within the XBP1s and PERK branches, but only in response to Tg and not Tm. Tg inhibits the sarcoplasmic/endoplasmic reticulum Ca^2^+ATPase (SERCA), depleting ER calcium stores and disrupting calcium-dependent chaperone function^46^. In contrast, Tm blocks N-linked glycosylation, leading to the buildup of unfolded glycoproteins in the ER^45^. These findings suggest that neurons are more sensitive to ER calcium depletion than to disturbances in glycoprotein folding, and that the UPR becomes asymmetrically impaired with age. Alternatively, the lack of global UPR activation in neurons upon exposure to ER stressors at 30 and 50 days, as was seen in glia, may indicate the neurons are in proteostasis collapse or highly regulate activation of the UPR with age. The selective preservation of the XBP1s branch with age has been associated with increased longevity and enhanced proteostasis, acting through cell-nonautonomous mechanisms that support systemic protein homeostasis^5,68^. To date, no UPR challenge studies have been specifically conducted in human neurons, highlighting the relevance of this co-culture model for understanding human-specific aging mechanisms in the nervous system.

Glial monocultures were phenotypically similar to co-culture glia; however, among proteins shared between the two culture conditions, the extent of downregulation differed markedly. This indicates that, while overall expression profiles may be comparable, the magnitude of proteomic changes is influenced by the culture context. Importantly, analysis of UPR machinery revealed that the age-related impairment in glial UPR signaling is cell-intrinsic and not dependent on neuron-glia crosstalk. This finding underscores a glia-specific vulnerability to proteostasis disruption that persists independently of neuronal influence.

One of the main limitations of our study is the duration of the aging timepoints. Without extended neuronal aging, it remains challenging to determine when glial health begins to decline to an extent that significantly impacts neuronal proteostasis. Future work using this *in vitro* model could incorporate single-cell proteomics to better resolve cellular heterogeneity, particularly within glial populations, as bulk proteomic analysis averages across diverse cell states, which may obscure cell-type or state-specific changes. Despite these limitations, our findings fundamentally reframe our understanding of brain aging by demonstrating that proteostasis decline is not uniform across cell types. Specifically, neurons maintained robust proteostasis machinery and metabolic function with age, while glial cells showed pronounced deterioration without compensatory stress responses. These results reveal important new insights into the heterogeneous nature of brain aging and suggest that differential cellular vulnerabilities may drive age-related neurodegeneration. Moving forward, this model can be adapted to investigate drug treatments, disease-specific contexts, or the dynamics of select proteins of interest. Our current study offers a foundational analysis pipeline and establishes baseline proteomic measurements for this system.

## Methods

### SH-SY5Y and N2a Monoculture and lysis conditions

SH-SY5Y cells, a human derived neuroblastoma cell line, were purchased from ATCC (CRL-2266). Neuro2a (N2a) cells, a mouse derived neuroblastoma cell line, were a gift from Keri Tallman (Vanderbilt University, Nashville, TN, USA). Both cell types were cultured in Dulbecco’s modified Eagle medium (DMEM) supplemented with 10% fetal bovine serum (FBS), 1% penicillin/streptomycin (P/S), and 1% glutamine (Q). Cells were incubated at 37 °C at 5% carbon dioxide (CO_2_). SH-SY5Y cells were split at 80% confluency, incubated with trypsin for 2-5 mins, with the trypsin removed prior to replating. For collection, cells were placed on ice and washed twice with cold phosphate buffered saline (PBS; 137 mM NaCl, 2.7 mM KCl, 10 mM Na_2_HPO_4_, 1.8 mM KH_2_PO_4_, adjust to pH 7.4). After washing, cells were lysed directly on the tissue culture plate with radioimmunoprecipitation assay (RIPA) buffer (50 mM Tris (pH 7.5), 150 mM NaCl, 0.1% SDS, 1% Triton X-100, 0.5% deoxycholate) with Roche complete EDTA-free protease inhibitor (Sigma) at 4 °C (rocking) for 30 minutes. Following lysis, the suspension was scraped into 1.5 ml centrifuge tubes and centrifuged at 16,000 g for 15 minutes to remove cellular debris. The lysate was then transferred to fresh tubes and the protein concentration was quantified using a bicinchoninic acid assay (BCA; ThermoFisher Scientific)^33^.

### Dual-species Set Ratio and UPR Activation Validation Experiments

To prepare dual species set ratio samples, untreated SH-SY5Y (human) and N2a (mouse) lysates were mixed at fixed ratios of 0:100, 25:75, 50:50, 75:25, and 100:0. Alternatively, to induce global UPR activation in SH-SY5Y cells, cells were treated with Tm (500 nM; Cayman Chemical) and Tg (500 nM; Cayman Chemical) for 16 hours. Additionally, SH-SY5Y cells treated with DMSO (0.1% v/v; Sigma) served as a control. Lysates from UPR-activated and control SH-SY5Y cells (treated with Tm, Tg, or DMSO) were mixed with untreated N2a lysate at a 1:4 ratio (20% SH-SY5Y to 80% N2a) (n=4). For both validation experiments, lysate was mixed prior to protein trypsin digestion.

### Western Blot Analysis (Global UPR Activated SH-SY5Y Cells)

To validate global UPR activated SH-SY5Y cells prior to LC-MS/MS analysis, 6x Laemmli buffer (12% SDS, 125 mM Tris, pH 6.8, 20% glycerol, bromophenol blue, 100 mM DDT) was added to 60 µg of protein in lysates per sample and heated for 5 min at 95 °C. Samples were then separated on an 12% SDS-PAGE gel (60 V for 20 min, 160 V for 80 min) and transferred to a 0.45 µm PVDF membrane (100 V for 80 min) for Western blotting. Prior to blotting, the blot was blocked with 5% (w/v) molecular-grade non-fat milk in TBST buffer (Tris-buffered saline with Tween 20; 50 mM Tris, 1.5 M NaCl, 0.1% Tween 20 (pH 7.4)) for 30 min at RT. Anti-KDEL (Enzo, ADI-SPA-827-F), anti-PDIA4 (ProteinTech, 14712−1-AP), anti-ASNS (ProteinTech, 14681-1-AP), and anti-GAPDH (GeneTex, GTX627408) antibodies were used to probe Western blots at a 1:1000 dilution in TBST blocking buffer (0.1% Tween, 5% BSA, 0.1% Sodium Azide).

### iN generation and sample collection

A. **Cell culture:** H1 human ES cells were maintained in mTeSR media and passaged using ReLeSR in Matrigel coated 6-well plates. Mouse glial cells were isolated from P1-P2 CD1 mouse pup forebrains and dissociated using Papain digestion and maintained in DMEM high glucose with 5% FBS and 1% P/S. Mouse glial cells were passaged 1-2 times using Trypsin before seeding on Matrigel coated 24-well plates. All cells were incubated at 37 °C with 5% CO_2_ and passaged at 80% confluency.
B. **Virus Generation:** Lentiviral vectors were generated using HEK-293T cells. Briefly, HEK cells were passaged one day before transfection in antibiotic-free media to reach 70-80% confluency at the time of transfection. HEK cells were transfected with lentiviral packaging plasmids pRSV-Rev, pMDLg/pRRE, pCMV-VSV-G and transgene expression constructs using Fugene6 transfection reagent and Opti-MEM. HEK cells were incubated with transfection complexes for 24 hours and then incubated in collection media and harvested after 36-48 hrs. Lentiviral supernatants were centrifuged at 2000 rpm for 15 minutes to remove cellular debris. Viral supernatants were aliquoted and stored at −80 °C.
C. **Generation of induced human neuron (iN) cells:** iN cells were generated as previously described^16,18^. On day 0, hES cells were dissociated using Accutase and plated as a single cell suspension in mTeSR with 10 µM Rho-associated protein kinase inhibitor (ROCKi), Ngn2 and rtTA lentivirus, and 8ug/mL polybrene. After 24 hours, the media was changed to induction media using DMEM/F12 containing 1X N2, non-essential amino acids, 1X B27 supplement, 2µg/mL doxycycline, BDNF, NT-3, and mLaminin (all 1µg/mL). On day 2, the induction media was changed to selection media made using the same reagents as induction media with the addition of 1µg/mL puromycin. Following 48 hours of selection, developing iNs were gently dissociated using Accutase and re-plated with mouse glial cells and maintained in neurobasal plus media containing 1X Glutamax, B27 supplement, 5% FBS, BDNF, NT-3, mLaminin, and doxycycline. Approximately 1e6 iN cells were plated into one well of a 24-well plate. Following re-plating, half of the neurobasal media was changed every 3 days. Supplements BDNF, NT-3, mLaminin, and doxycycline were maintained in neurobasal media until day 15. After day 15, half of the neurobasal media was changed every 5 days without the addition of neurotrophic supplements and doxycycline.
D. **Treatment of Co-cultures with Global UPR Activators (Tm & Tg):** Co-cultures were treated at days 30 and 50 for 16 hours with Tm (5 µM; Cayman Chemicals) and Tg (300 nM; Cayman Chemicals), prior to collection. Treatment concentrations with UPR activators was determined in a previous study^34^.
E. **Sample collection:** Culture media was first aspirated from the culture well and samples were rinsed with sterile PBS. Samples were then collected in a buffer containing 1X RIPA lysis buffer, 1X protease, and 1X phosphatase inhibitors dissolved in water. 75µL of collection buffer was added to each well of a 24-well plate. Cells were then mechanically dissociated using a P200 pipette tip containing the collection buffer. Samples were placed in a collection tube and immediately placed on ice prior to storage at −80 °C. Thawed lysates were centrifuged at 16,000 g for 15 minutes to remove cellular debris. Cleared lysate was then normalized with a BCA prior to digestion.

### Protein Digestion and Sample Preparation for LC-MS/MS

Protein cleanup and digestion was carried out on the Biomek i5 (Beckman Coulter). For experiments with SH-SY5Y and N2a lysate, a total of 20 µg of protein was digested. Alternatively, 10 µg of protein from each co-culture or glia mono-culture sample was aliquoted for digestion using the automated sample handler. Proteins were reduced with fresh dithiothreitol (DTT; 5 mM, Sigma) for 30 min at 37 °C/ 1000 rpm and alkylated with fresh iodoacetamide (20 mM, Sigma) for 15 min at RT (in darkness). To quench the alkylation, DDT was added a second time (5 mM) with incubation for 15 min. Subsequently, proteins were digested on magnetic single pot solid phase enhanced sample preparation beads (SP3; Cytiva; 50 mg/mL; 1:1 hydrophilic to hydrophobic beads) and cleaned-up as previously described^35^. Protein digestion was performed using 10 and 20 µg of protein with 4 and 8 µL of SP3 beads, respectively. Protein was bound to SP3 beads with 100% ethanol and washed three times with 80% ethanol for clean-up. Proteins were then digested with Trypsin/Lys-C (Thermo Fisher; 1:40 protease to protein ratio) in 50 µL ammonium bicarbonate (ABC; 100 mM; pH 8) at 37 °C / 700 rpm for 10 hours. Digested peptides were removed from SP3 beads after digestion on the Biomek i5 and dispensed manually into low-bind tubes. Formic acid (FA) was added to each sample to a final concentration of 2% (v/v) and samples were subsequently dried *in vacuo* using a SpeedVac (ThermoFisher Scientific, SPD111V). Samples were stored dried at −80 °C until use. Peptides were resuspended to a final concentration of 200 ng/µL in buffer A (4.9% acetonitrile, 95% H_2_O, 0.1% FA (v/v/v). Samples were then spun down at 21,100 g for 15 min and transferred to a fresh low bind tube prior to LC-MS/MS analysis.

### LC-MS/MS Analysis

LC-MS/MS analysis was performed using an Exploris480 mass spectrometer (Thermo Fisher) equipped with a Dionex Ultimate 3000 RSLCnano system (Thermo Fisher). Peptides were separated using a 21.5 cm fused silica microcapillary column (ID 100 µm) ending with a laser-pulled tip filled with Aqua C18, 3 µm, 100 Å resin (Phenomenex # 04A-4311). Electrospray ionization was performed directly from the analytical column by applying a voltage of 2.2 kV (positive ionization mode) with an MS inlet capillary temperature of 275 °C and a RF Lens of 40%.

A total of 600 ng of peptides per sample were loaded onto a commercial trap column (C18, 5 µm, 0.3×5mm; ThermoFisher Scientific, 160454) using an autosampler. Peptides were then eluted and separated on a 2 h gradient with a constant flow rate of 500 nL/min: 2% B (5 min hold) was ramped to 35% B over 90 min and stepped to 80% in 5 min and held at 80% B for 5 min, followed by a 13 min hold at 4% B to re-equilibrate the column. Between injections, a 45 min column-wash was performed using the following gradient: 2% B (6 min hold) stepped to 5% over 2 min and subsequently ramped to a mobile phase concentration of 35% B over 7 min, ramped to 65% B over 5 min, held at 85% B for 8 min, then returned to 3% B for the remainder of the analysis.

The DIA-MS method consisted of 62 MS/MS scan events with a 10 *m/z* precursor isolation window and 1 *m/z* overlap (30,000 resolution, 1e7 normalized ACG target, 55 ms maxIT, 27% NCE, 40% RF lens) taken across a 380-1000 *m/z* mass range. A precursor spectrum (380-1000 *m/z*, 3e6 normalized ACG target, 40% RF lens) was taken at 120,000 resolution with a maxIT of 240 ms. Loop control was utilized with n= 20 to intersperse MS^1^ scans between MS/MS acquisitions. The 66 spectra collected per cycle resulted in a duty cycle time of approximately 4.4 sec. All DIA data was collected in profile and positive mode. Alternatively, for GPF analysis of dual-species set ratio and aged co-culture samples, five separate DIA methods were generated with precursor mass ranges of: 380-504, 504-628, 628-752, 752-876, and 876-1000 *m/z*. For each method, the isolation window was set at 2 *m/z* with a 1 *m/z* overlap. All other parameters outlined above were kept constant.^36^

### Peptide Identification and Quantification

All DIA spectra were analyzed using DIA-NN (version 1.8.1) with a library free search workflow. For GPF experiments, an initial spectral library was generated in DIA-NN using GPF LC-MS/MS files along with the corresponding sample files. This library was then used in a second DIA-NN analysis to obtain peptide quantifications for the samples of interest. Spectral libraries were generated using a Uniprot SwissProt canonical human (downloaded 01May2024; containing 20,434 entries) and mouse (downloaded 01May2024; containing 17,214 entries) FASTA database and contaminant FASTA^37^ (containing 379 entries). Peptides ranging from 7 to 30 amino acids in length were considered, allowing for one missed trypsin cleavage and a maximum of one variable modification (Met oxidation or N-terminal acetylation). N-terminal M excision was enabled and cysteine carbamidomethylation was included as a fixed modification. Precursors between *m/z* 380-1000 with charge states of 1-4 were considered. Additionally, fragment ions between *m/z* 200 and 1800 were included. Deep learning (using a single-pass mode neural network classifier) was then utilized to generate a new *in-silico* spectral library from the DIA data provided to then research the raw files against. Smart profiling was used for library generation. A mass accuracy tolerance of 10 ppm for precursor ions (MS^1^) and 15 ppm for fragment ions (MS^2^) was applied during the database search, with a 1% FDR at the peptide and protein levels. Protein inference was made on genes and robust LC (high precision) was used as the quantification strategy. Protein assembly was performed in DIA-NN based on the parsimony principle.

### Data Analysis (Proteomics)

A custom R script was developed to process DIA-NN output reports to perform statistical filtering, species-specific peptide assignment, normalization, data visualization, and UPR target filtering. As part of this workflow, the diann.dia.qc() function was used to retain peptides with a peptide *q*-value, global protein *q*-value, and posterior error probability (PEP) score below 1%. To increase confidence in protein identifications, only proteins supported by at least two peptides were retained for downstream analysis. Contaminant proteins^37^ were removed prior to subsequent processing.

To classify LC-MS/MS detected peptides species-specifically as human, mouse, or shared (i.e., present in both human and mouse proteins), a theoretical peptide database was first generated using the FASTA files input into DIA-NN. Proteins from these FASTA files were digested *in silico* into tryptic peptides, allowing zero or one missed cleavages and peptide lengths between 6 and 31 amino acids, using the cleaver R package via our custom digest.two.species() function. Next, empirical peptides were matched to the theoretical peptide database to assign species origin using the match.data.sm() function. Finally, peptides that remained unclassified were searched against the original FASTA files using the rematch() function to determine their species of origin.

Following peptide-level filtering and species-specific peptide assignment, data was globally median normalized using the medNorm() function. For each sample *i*, a normalization factor F_*i*_ was computed as the ratio between the global median of all precursor intensities (M_G) and the median intensity within a particular sample (M_I). The normalized abundance 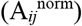 for each precursor *j* in sample *i* (A_*ij*_) was then calculated in the following two steps: (1)F_*i*_ = M_G / M_I & 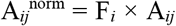. Protein-level quantifications were then inferred from the filtered and normalized peptides using the max_LFQ() function from the DIA-NN R package^38^. After log_2_ transformation of the resulting protein intensity matrices, data were filtered using the filter.NA() function to ensure robustness of protein-level quantification. For the aging co-culture dataset (n = 10 per timepoint), a minimum of three valid observations per timepoint was required. No imputation was applied.

To assess the average expression change for each treatment condition, the mean log_2_ fold-change (log_2_FC) across biological replicates was computed for each protein using the average.FC() function in R. First, the control mean (µ_i_) was calculated using the log_2_ intensity values, *x*, of each protein, *i*, where *k* is the number of control samples, as shown in Equation (1). Then, the log_2_ fold change for each sample *j* (including both control and treatment samples) was computed using Equation (2). Finally, the average for each condition was calculated using the base R function rowMeans().

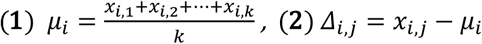

Identified proteins were subsequently filtered against branch-specific UPR targets to evaluate pathway-specific activation. All volcano plots were generated in PRISM using unpaired t-tests with multiple testing corrections. Proteins were deemed significant if they had a −log_10_(adjusted p-value) greater than 1.3, corresponding to a 5% false discovery rate (FDR). The FDR was controlled using the two-stage step-up method of Benjamini, Krieger, and Yekutieli. Ontology analysis of differentially regulated proteins was performed using Enrichr^39–41^, with an adjusted p-value of less than 0.05 considered statistically significant.

### Experimental Design and Statistical Rationale

For validation of the developed pipeline, a dual-species set ratio experiment was conducted using a total of 20 samples. Four biological replicates were prepared for each untreated human lysate (SH-SY5Y) percentage (0%, 25%, 50%, 75%, and 100%), balanced with untreated mouse lysate (N2a). Secondly, a total of 12 samples were generated to evaluate the detection of UPR activation in dual-species compared to single-species samples. Four biological replicates were prepared for each of the three drug treatments: Tunicamycin (Tm), Thapsigargin (Tg), and dimethyl sulfoxide (DMSO). A total of 40 aged co-culture samples were generated, with cells collected at 30, 40, 50, and 60 days, and 10 biological replicates per timepoint. In addition, 20 aged glia monoculture samples were prepared, with cells collected at the same timepoints and 5 biological replicates per timepoint. For the co-culture UPR challenge experiments, 24 samples were generated by treating co-cultures at days 30 and 50 with either Tm or Tg, with 7 biological replicates per condition. To ensure quality of digestion on the Biomek i5, 20 µg of *E. coli* lysate was digested and analyzed via DIA LC-MS/MS. Additionally, prior to the start of each MS analysis, 600 ng of *E. coli* peptides were measured with unfractionated DIA to ensure consistency of instrument performance. Statistical tests used to process proteomics data can be found in the methods section titled: “Data Analysis (Proteomics).”

## Data, Materials, and Software Availability

Mass spectrometry data have been deposited to ProteomeXchange Consortium via the PRIDE partner repository with the project accession (PXD066290) and project DOI (10.6019/PXD066290).

## Acknowledgements

This work was supported by R35 GM133552 (L.P) and R01 NS134128 (E.K.).

## Author Contributions

Conceptualization, L.A.B., B.U., L.P, E.K.; Investigation, L.A.B., S.K.G., A.H., B.U., M.K.; Writing – Original Draft, L.A.B.; Writing – Review and Editing, L.A.B., A.H., E.K., M.K.; L.P.; Visualization, L.A.B.; Supervision, E.K., L.P.; Funding Acquisition, E.K., L.P.

## Supporting Information

**SI - PDF**

**Dataset S1**. Number of human and mouse false positive identifications depending on statistical filtering conditions (1 or 2).

**Dataset S2.** Number of peptide/ protein identifications and summed abundances of identified proteins whether the data was searched in DIA-NN with library-free mode or with a library (gas-phase fractionation generated).

**Dataset S3**. Co-culture human and mouse proteins (Log_2_ protein abundance, ≥ 2 peptides per protein, filtered at three observations per protein per timepoint).

**Dataset S4**. Human and Mouse UPR Proteomics Targets.

**Dataset S5**. UPR activated co-culture human and mouse proteins (Log_2_ protein abundance, ≥ 2 peptides per protein, filtered at three observations per protein per timepoint).

**Dataset S6**. Glia monoculture (Log_2_ protein abundance, ≥ 2 peptides per protein, filtered at three observations per protein per timepoint).

