## Supplemental Information for "Proteostasis and Unfolded Protein Response Dynamics in Human Neuron and Mouse Glia Co-culture Reveal Cell-Specific Aging Responses"

**Table of Contents**

Figures S1-S14…...……………………………………………………………………...…..2-14

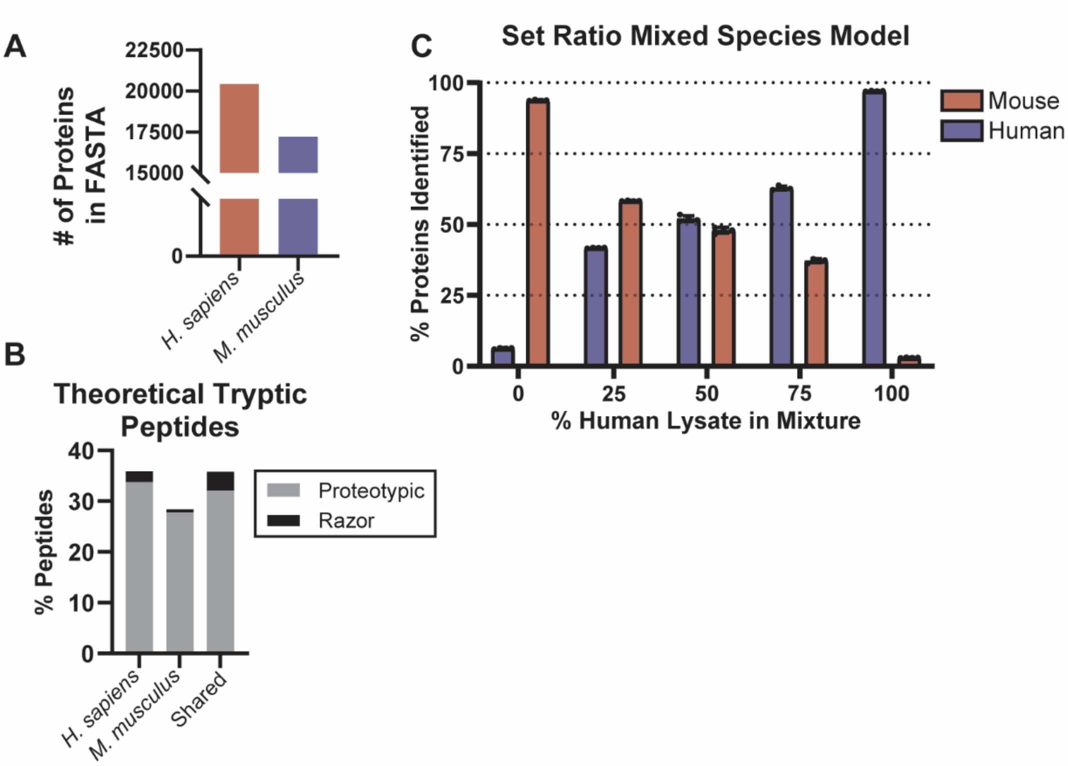


**Figure S1. Development and validation of a pipeline to analyze dual-species co-culture samples via DIA LC-MS/MS. (A)** Number of protein entries from human and mouse FASTA files used to construct the theoretical tryptic peptide database. **(B)** The percentage of peptides in the theoretical tryptic database, generated using the R package *cleaver*, that are classified as proteotypic or razor, and categorized as human, mouse, or shared. **(C)** Number of human or mouse proteins identified in samples with increasing amounts of human lysate to mouse lysate and vice versa (n = 4).

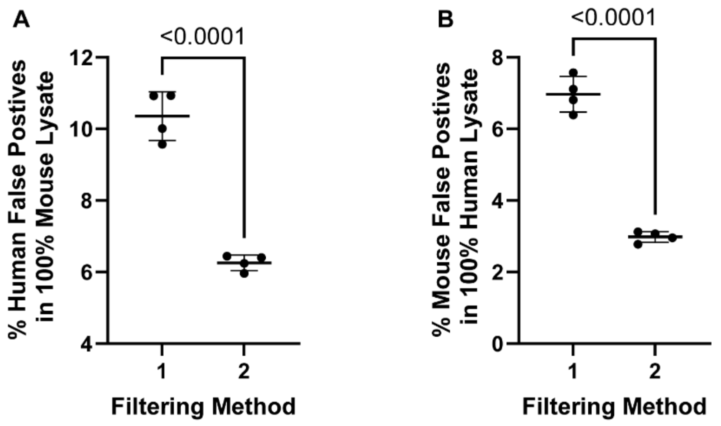


**Figure S2. Comparison of two filtering methods (method 1 without PEP score and method 2 with PEP) applied in R-studio to the DIA-NN output files for the analysis of dual-species co-culture samples. (A)** The percentage of human false positive protein identifications in the 100% mouse lysate samples and **(B)** the percentage of mouse false positive protein identifications in the 100% human lysate samples (n = 4). Shown are the mean ± SD (n = 4).

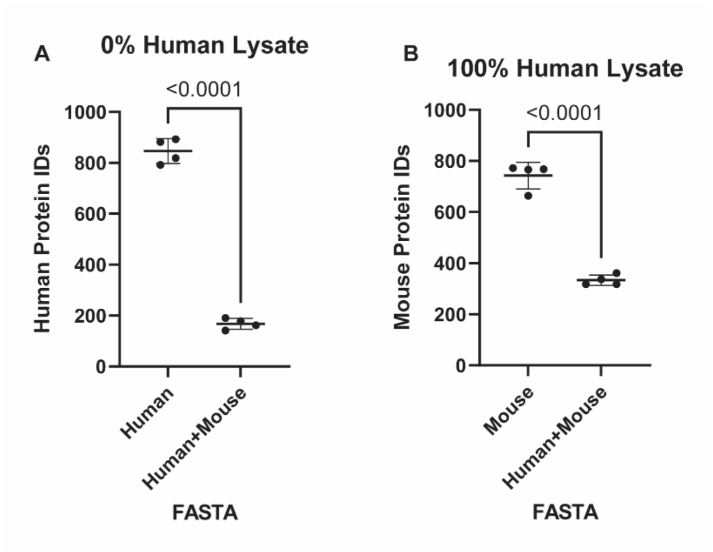


**Figure S3. False positive protein identifications introduced by single species and multi-species FASTA database searching. (A)** Number of human proteins identified (≥ 2 peptides per protein at 1% FDR) when analyzing a 100% mouse lysate sample in DIA-NN using a human FASTA and a combined human/mouse FASTA, with species-specific peptide assignment in R. **(B)** Number of mouse proteins identified (≥ 2 peptides per protein at 1% FDR) when analyzing a 100% human lysate sample using a mouse FASTA and a combined human/mouse FASTA, with species-specific peptide assignment in R. Shown are the mean ± SD (n = 4).

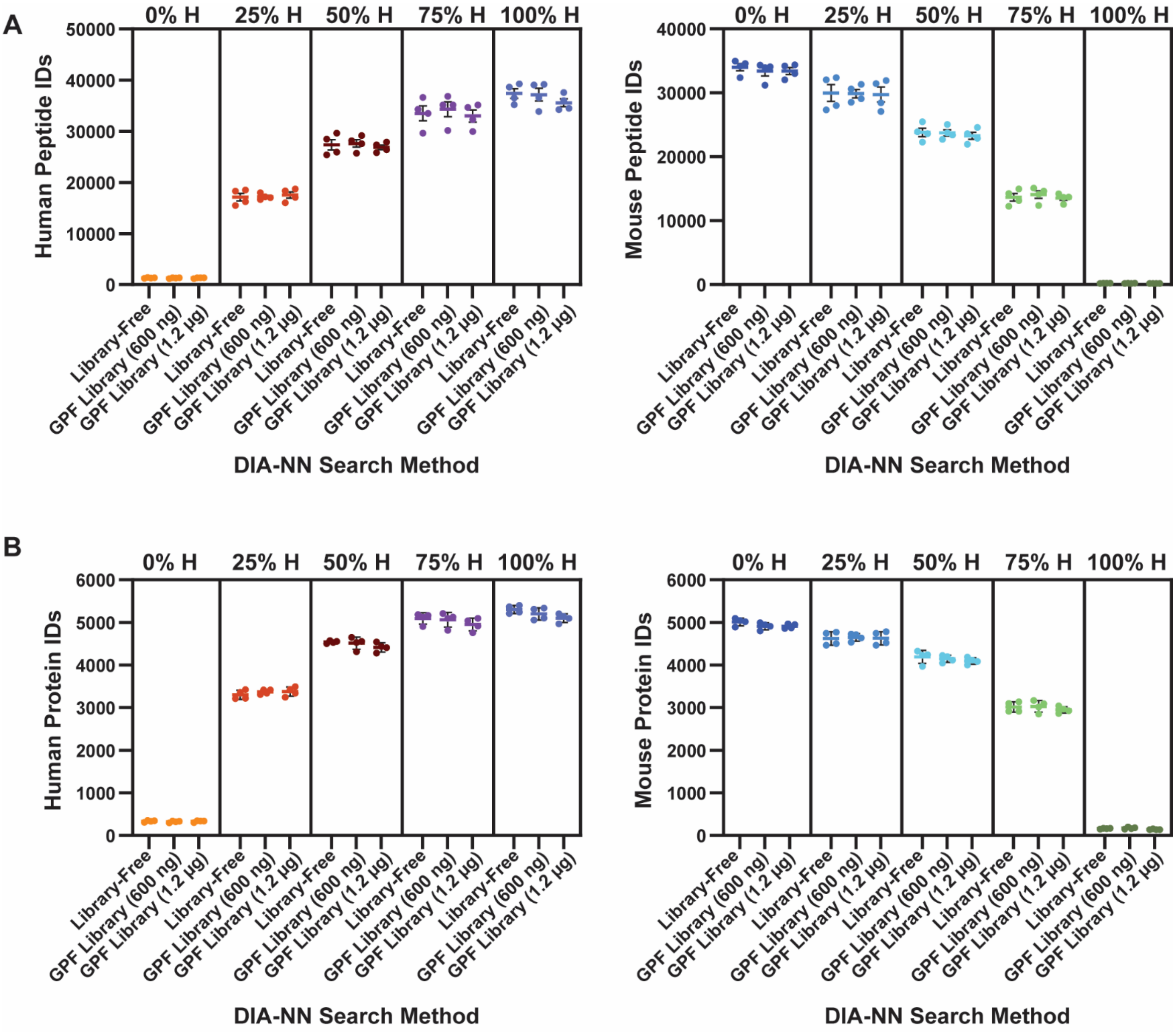


**Figure S4. Species-specific peptide and protein identification rates did not significantly increase when using DIA-NN with a sample-specific spectral library generated via gas-phase fractionation (GPF) compared to library-free analysis.** Number of peptides (**A**) and proteins (**B**) identified in mixed species samples when utilizing library-free and GPF spectral library modes in DIA-NN. Error bars represent the mean ± SD.

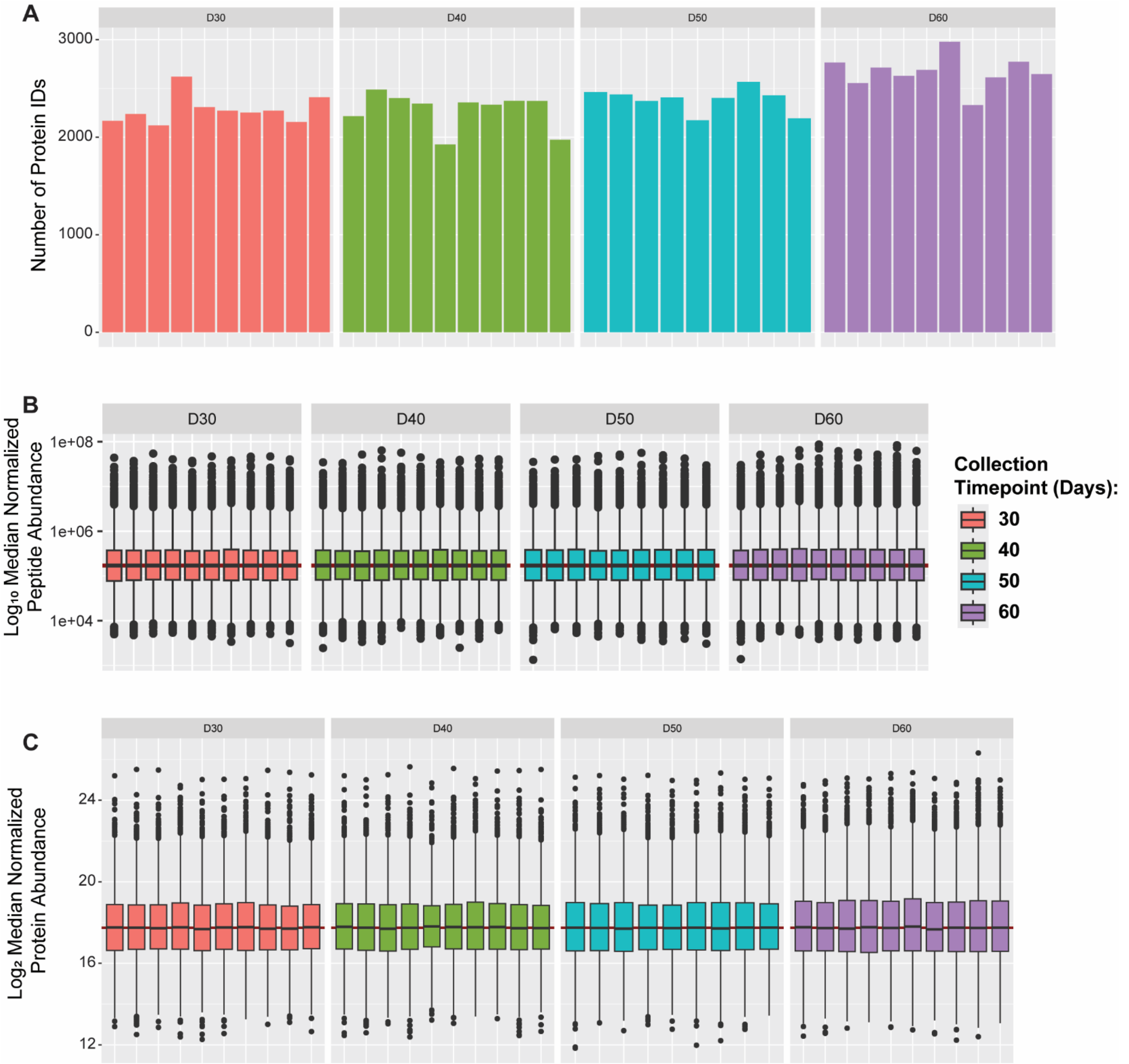


**Figure S5. Validation of DIA-MS analysis, human neuronal proteins identified in aged co-culture samples. (A)** Number of neuronal proteins (human) identified in co-cultures via DIA-MS across injections, separated by collection timepoint. Box and whisker plots show the distribution of median normalized **(B)** peptide abundances and **(C)** protein abundances separated by collection timepoint (n = 10). Box and whisker plots represent all peptide or protein abundances. The top and bottom of the box represent the third (Q3) and first (Q1) quartiles, respectively, with the middle line indicating the median. The whiskers extend to the smallest and largest values within 1.5 times the interquartile range (IQR).

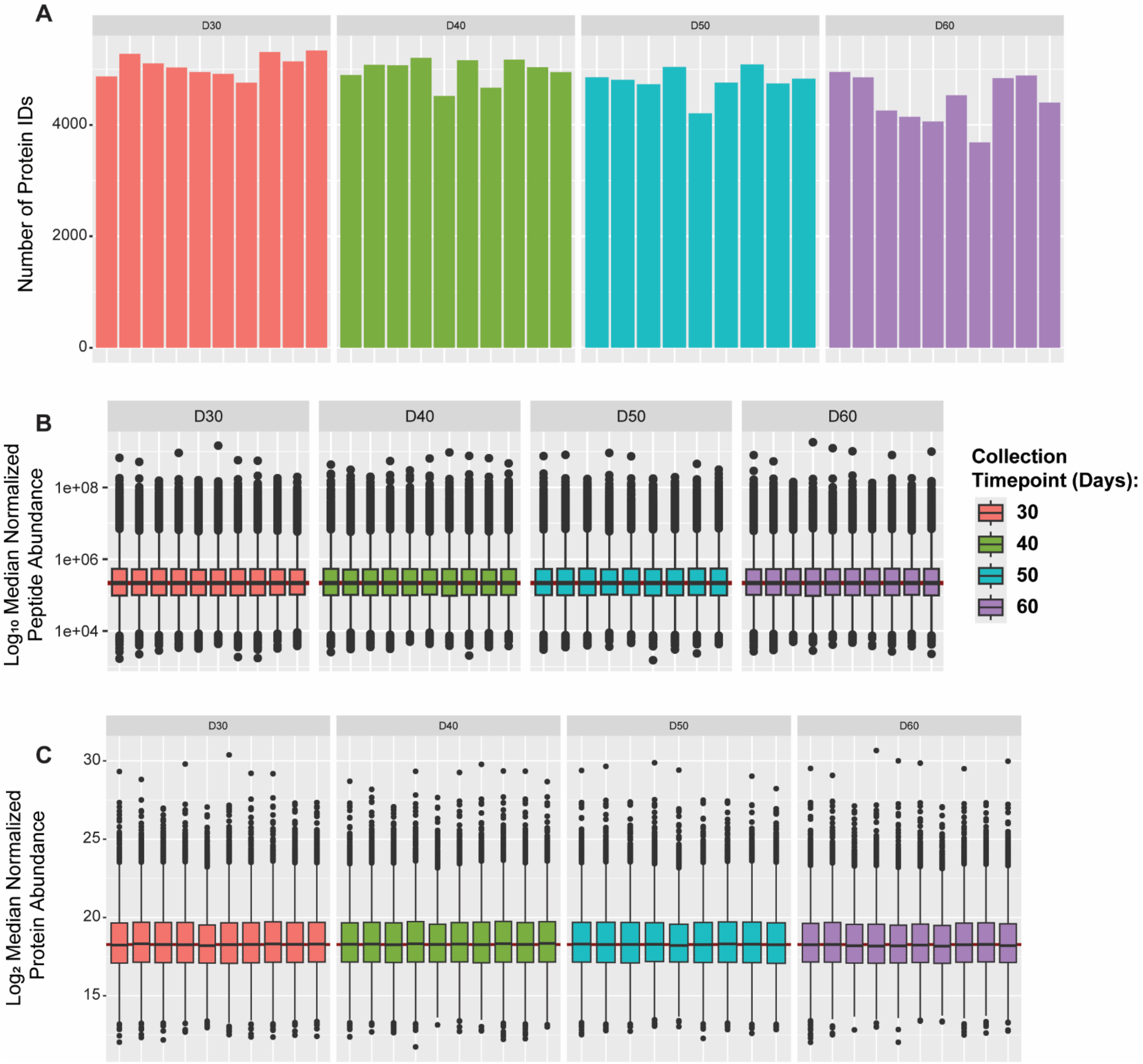


**Figure S6. Validation of DIA-MS analysis, glial mouse proteins identified in aged co-culture samples. (A)** Number of glial proteins identified in co-cultures via DIA-MS across injections, separated by collection timepoint. Box and whisker plots show the distribution of median normalized **(B)** peptide abundances and **(C)** protein abundances separated by collection timepoint (n = 10). Box and whisker plots represent all peptide or protein abundances. The top and bottom of the box represents the third (Q3) and first (Q1) quartiles, respectively, with the middle line indicating the median. The whiskers extend to the smallest and largest values within 1.5 times the interquartile range (IQR).

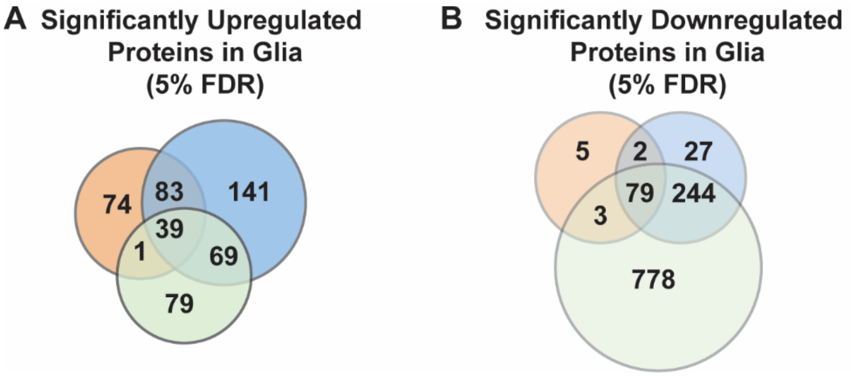


**Figure S7. Evaluation of significantly upregulated and downregulated glial mouse proteins identified in aging co-culture samples.** Significantly upregulated (**A**) and downregulated (**B**) proteins (5% FDR) shared and unique to the 40 (orange), 50 (blue), and 60 (green) day collection timepoints.

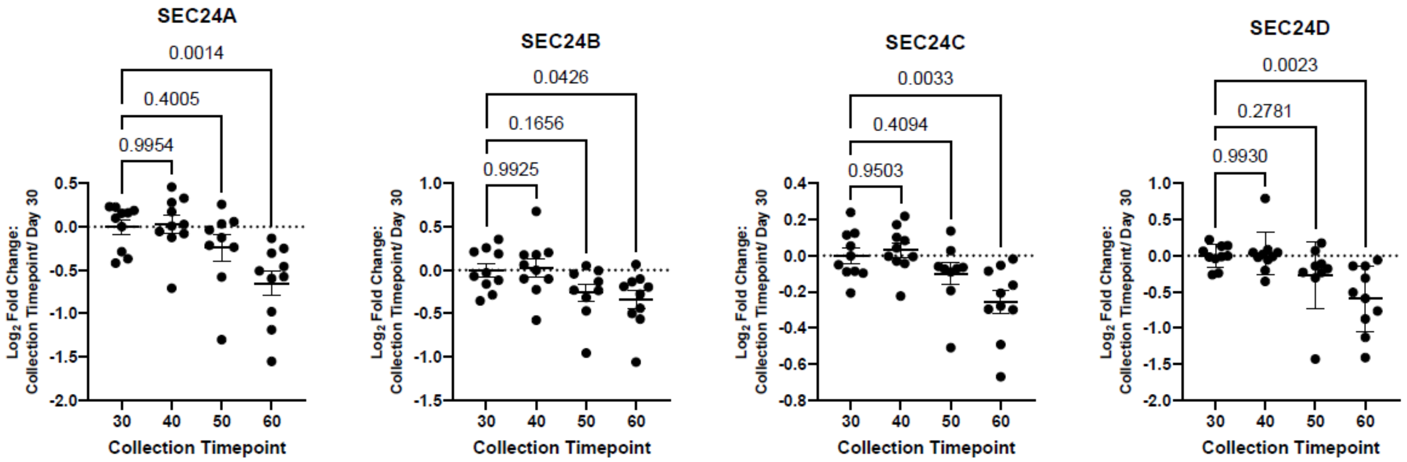


**Figure S8. Quantification of SEC24 isoforms in neurons at different ages (30, 40, 50, and 60 days) using LC-MS/MS**. $Log_2$ fold changes at each timepoint are shown relative to the day 30 control. n = 10; data are presented as mean ± SEM. Statistical analysis was performed using one-way ANOVA, followed by Dunnett's post hoc test for multiple comparisons against the day 30 control. $p < 0.05$ was considered statistically significant. Error bars represent the mean ± standard error of the mean (SEM).

.

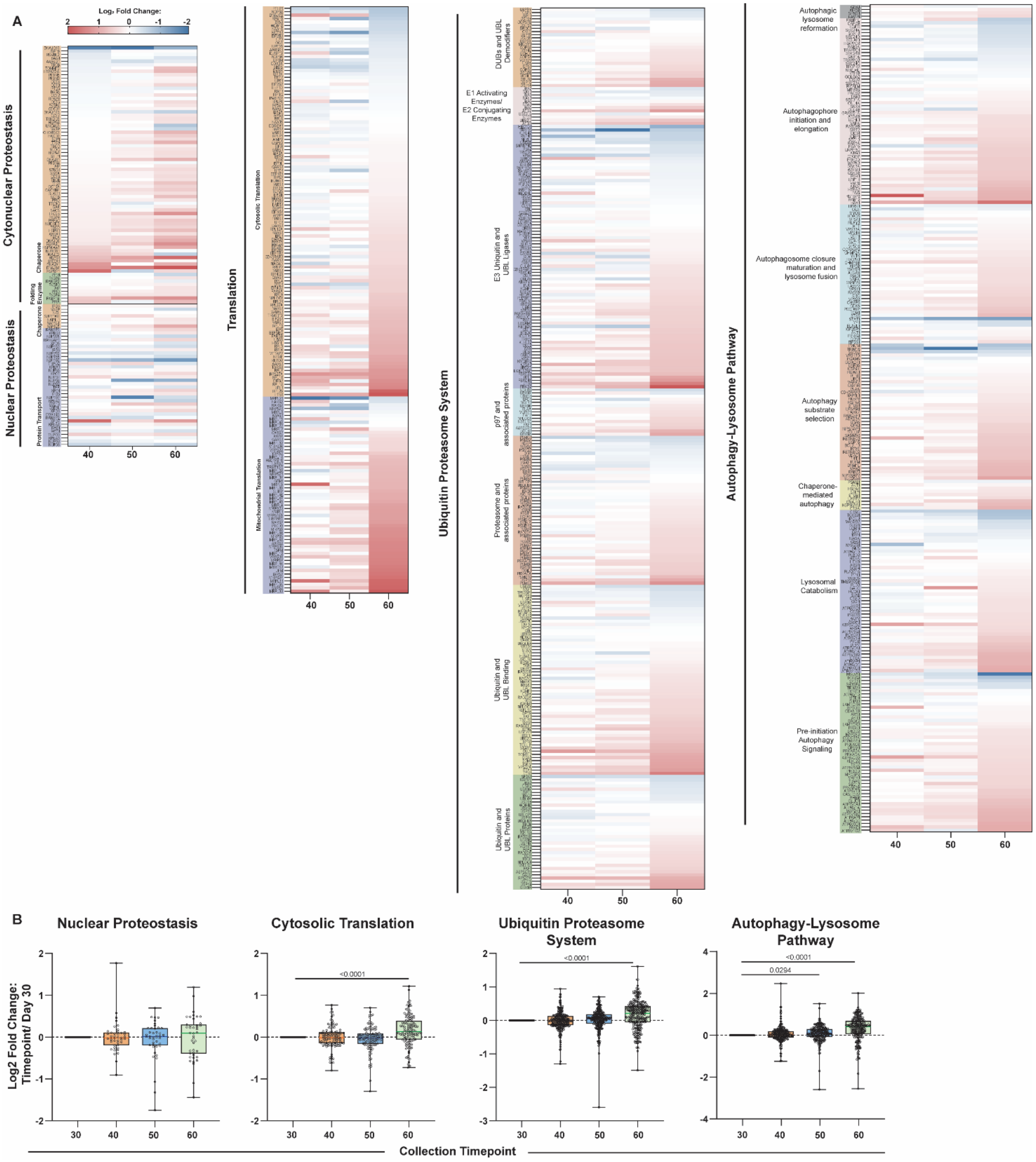


**Figure S9. Heatmaps of human neuronal proteins involved in proteostasis.** **(A)** Heatmap of the $\log_2$ fold change of neuronal proteins involved in cytonuclear and nuclear proteostasis, translation, the ubiquitin proteosome system, and autophagy-lysosome pathway identified at days 40, 50, and 60 relative to the day 30 control. Proteins were annotated and categorized by the Proteostasis Consortium (cite). **(B)** Boxplots showing the $\log_2$ fold change (collection timepoint/day 30) of neuronal proteins involved in nuclear proteostasis, cytosolic translation, the ubiquitin proteosome system, and autophagy-lysosome pathway, plotted collectively. n = 10; data are presented as mean ± SEM. Statistical analysis was performed using one-way ANOVA, followed by Dunnett's post hoc test for multiple comparisons against the day 30 control. $p < 0.05$ was considered statistically significant.

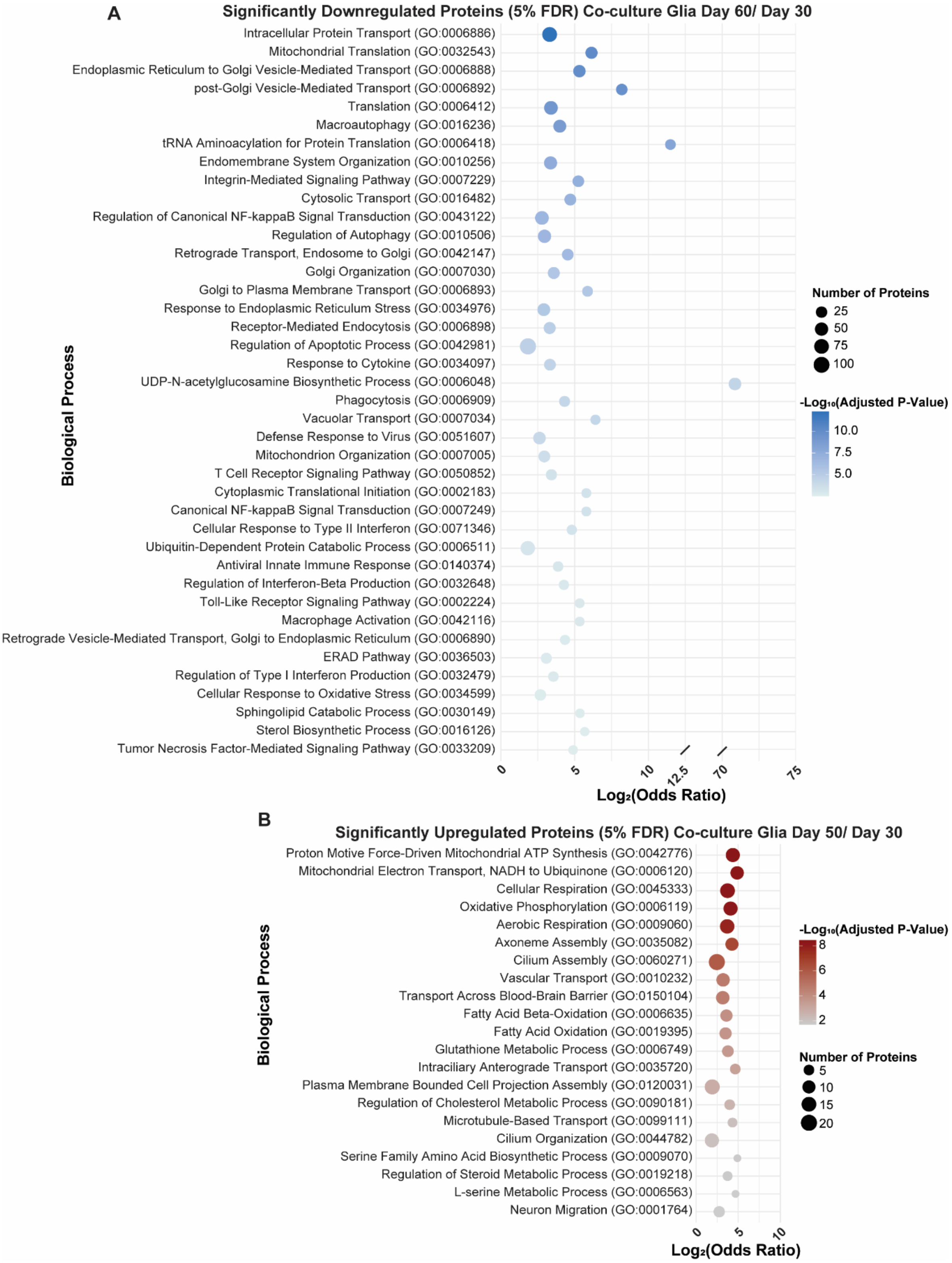


**Figure S10. Evaluation of significantly upregulated and downregulated glial mouse proteins identified in aging co-culture samples.** Gene ontology analysis of significantly downregulated glial proteins at day 60 compared to day 30 (**A**) and upregulated glial proteins (Day 50/ Day 30; **B**) and in the co-culture performed using Enrichr[1]. Adjusted p-value was computed using the Benjamini-Hochberg method to correct multiple hypothesis testing.

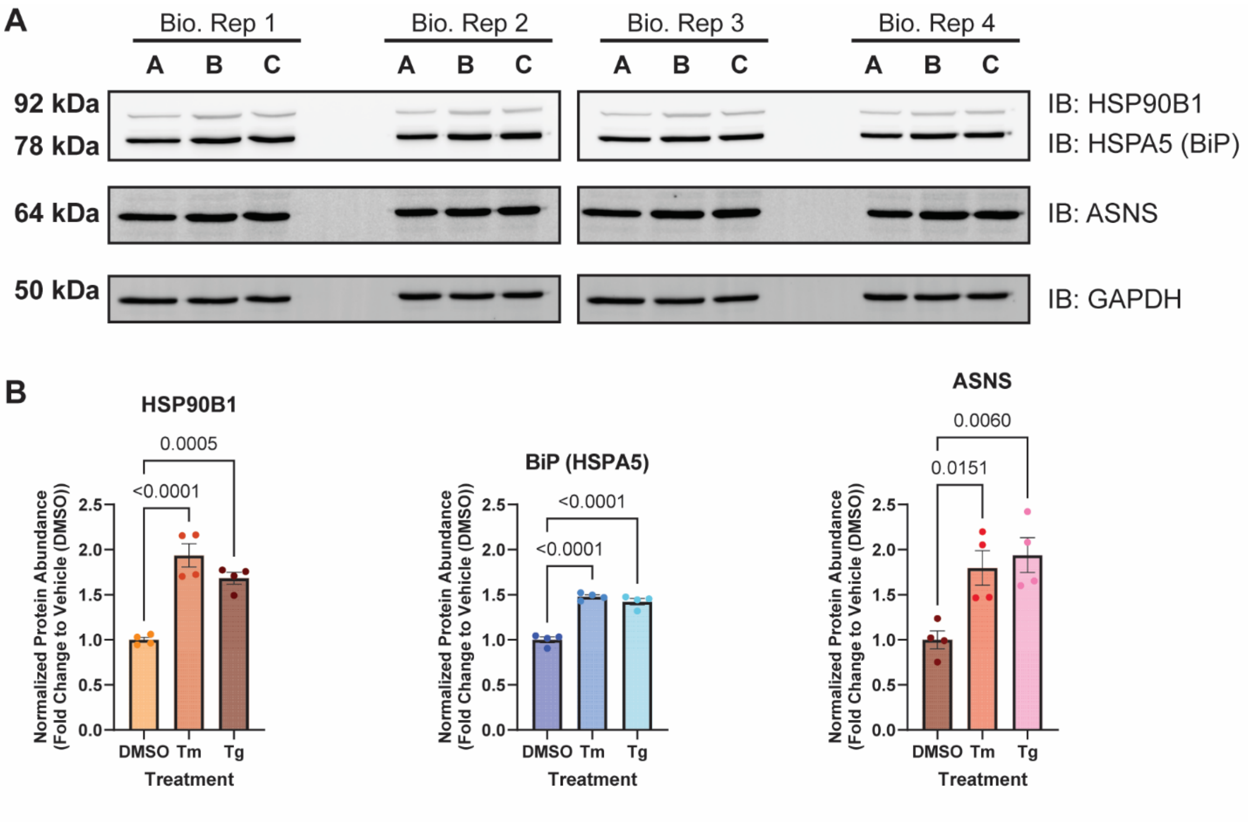


**Figure S11. Validation of UPR activation in SH-SY5Y cells with quantitative western blot.** (**A**) Western blots of UPR protein markers (HSP90B1 (GRP94), BiP (HSPA5/GRP78), and ASNS) in HEK293 cells treated with DMSO (0.1%), Tm (500 nM), and Tg (500 nM) for 16 h. GAPDH was used as a housekeeping gene for loading control. Displayed blot sections are from the same blot image and exposure settings. (**B**) Quantification of western blots in panel (A), normalized to GAPDH band intensities and compared to the DMSO control (vehicle). n = 4, bars represent the mean and errors represent ± SEM. Statistical analysis was performed using a one-way ANOVA with Benjamini, Krieger, and Yekutieli multiple testing correction, $p < 0.05$ considered statistically significant.

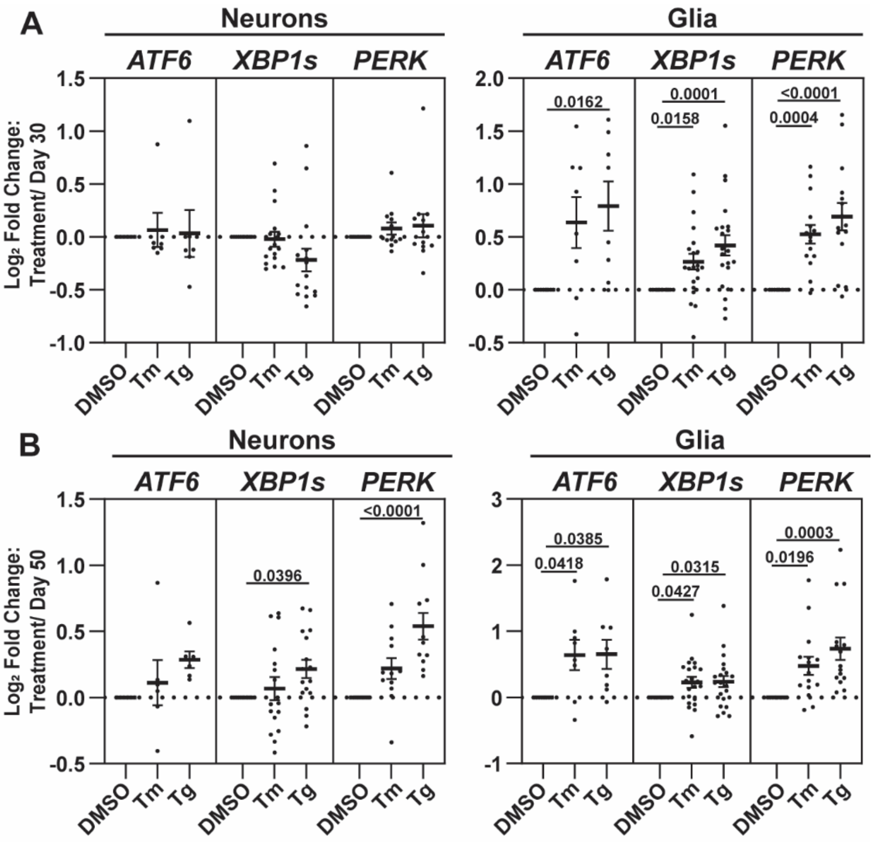


**Figure S12. Treatment of co-culture samples at days 30 and 50 with DMSO (Vehicle), Tm, and Tg for 16 hours.** (**A** and **B**) n = 7, mean ± SEM. Statistical analysis was performed using a one-way ANOVA test with Dunnett's post hoc test was computed. A $p$ value < 0.05 was considered statistically significant.

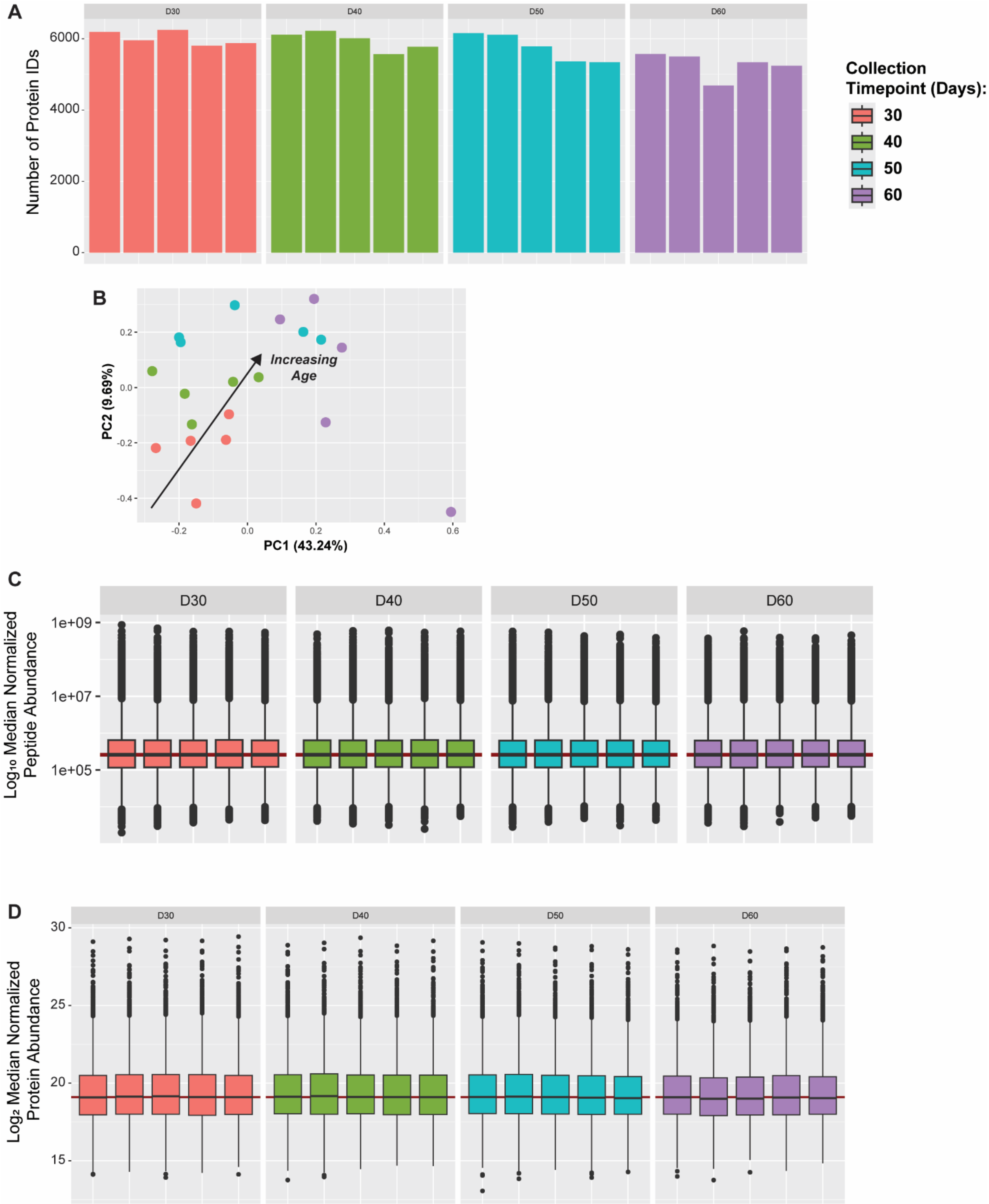


**Figure S13. Validation of DIA-MS analysis, glial proteins identified in monocultures. (A)** Number of mouse proteins identified via DIA-MS across injections, separated by collection timepoint. Box and whisker plots showing the distribution of median normalized **(B)** peptide abundances and **(C)** protein abundances separated by collection timepoint (n = 10). Box and whisker plots represent all peptide or protein abundances. The top and bottom of the box represent the third (Q3) and first (Q1) quartiles, respectively, with the middle line indicating the median. The whiskers extend to the smallest and largest values within 1.5 times the interquartile range (IQR)

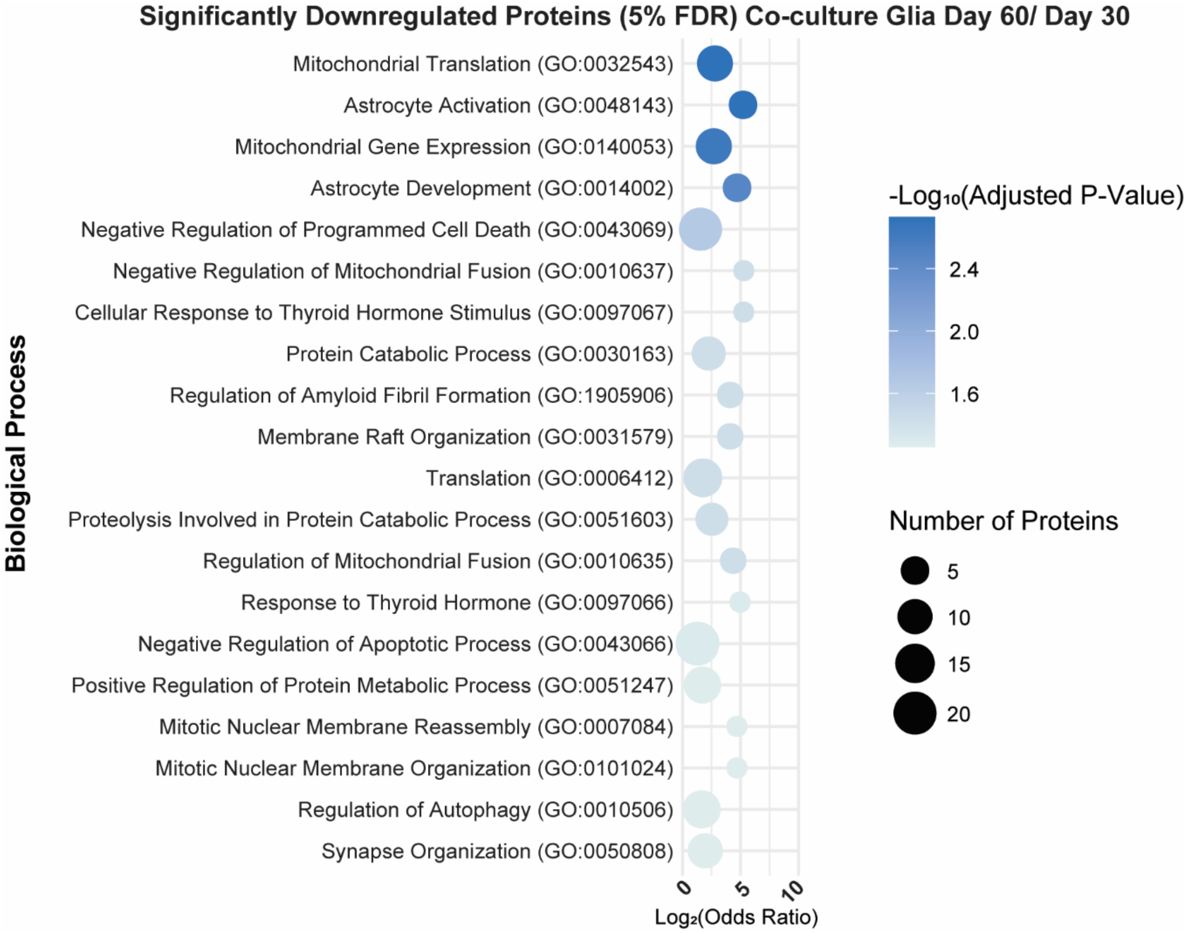


**Figure S14. Ontology of significantly downregulated proteins unique to co-culture glia.** Significantly downregulated biological processes associated with proteins unique to co-culture glia, as opposed to those shared with or unique to monoculture glia, identified using Enrichr. Adjusted p-value was computed using the Benjamini-Hochberg method to correct multiple hypothesis testing.
